## Supplemental Figures and Tables for "A chromosome-level genome sequence of a model chrysanthemum: evolution and reference for hexaploid cultivated chrysanthemum"

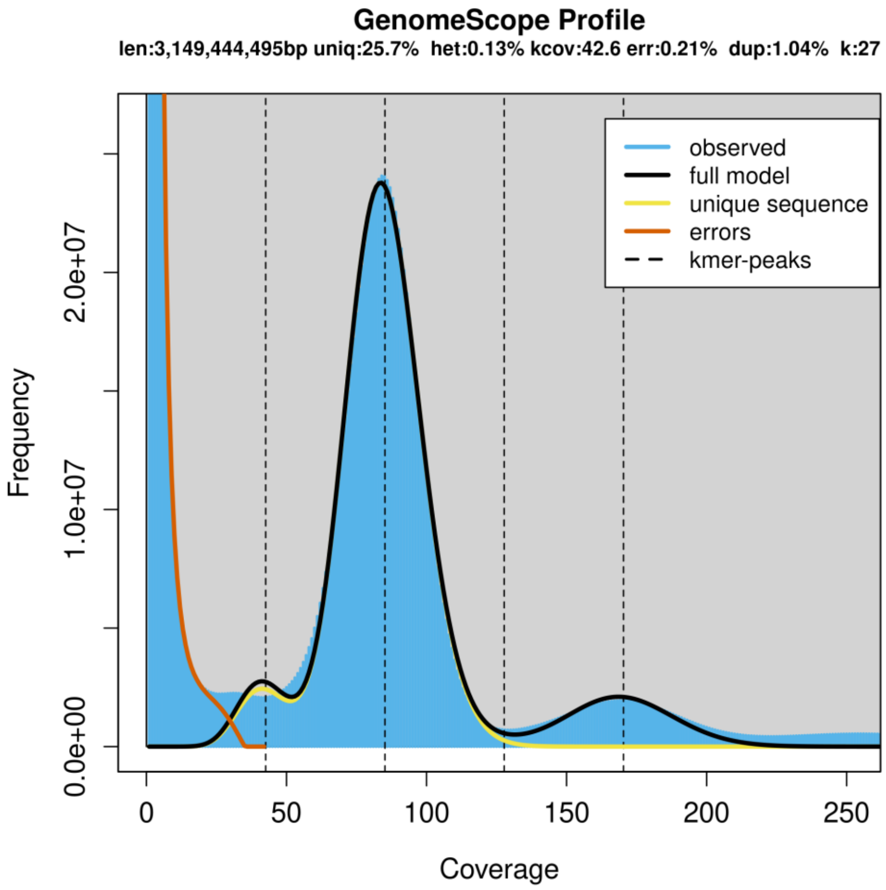


**Supplementary Figure 1 | *k*-mer analysis of unassembled short-read sequences.**

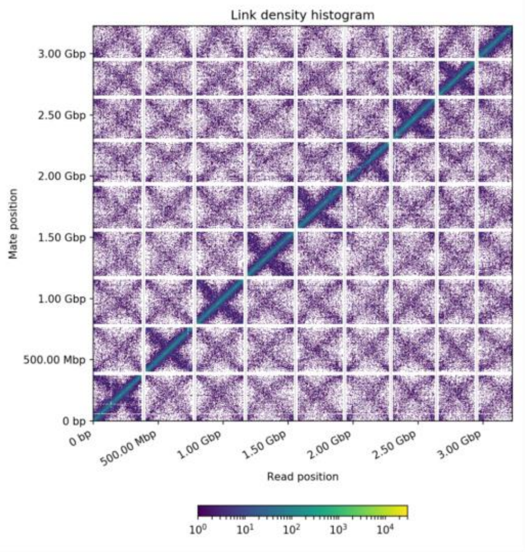


**Supplementary Figure 2 | Heat map of Hi-C interactions among the nine largest scaffolds of Gojo-0 genome.** The link density of paired sequencing reads in different contigs is illustrated by a color gradient.

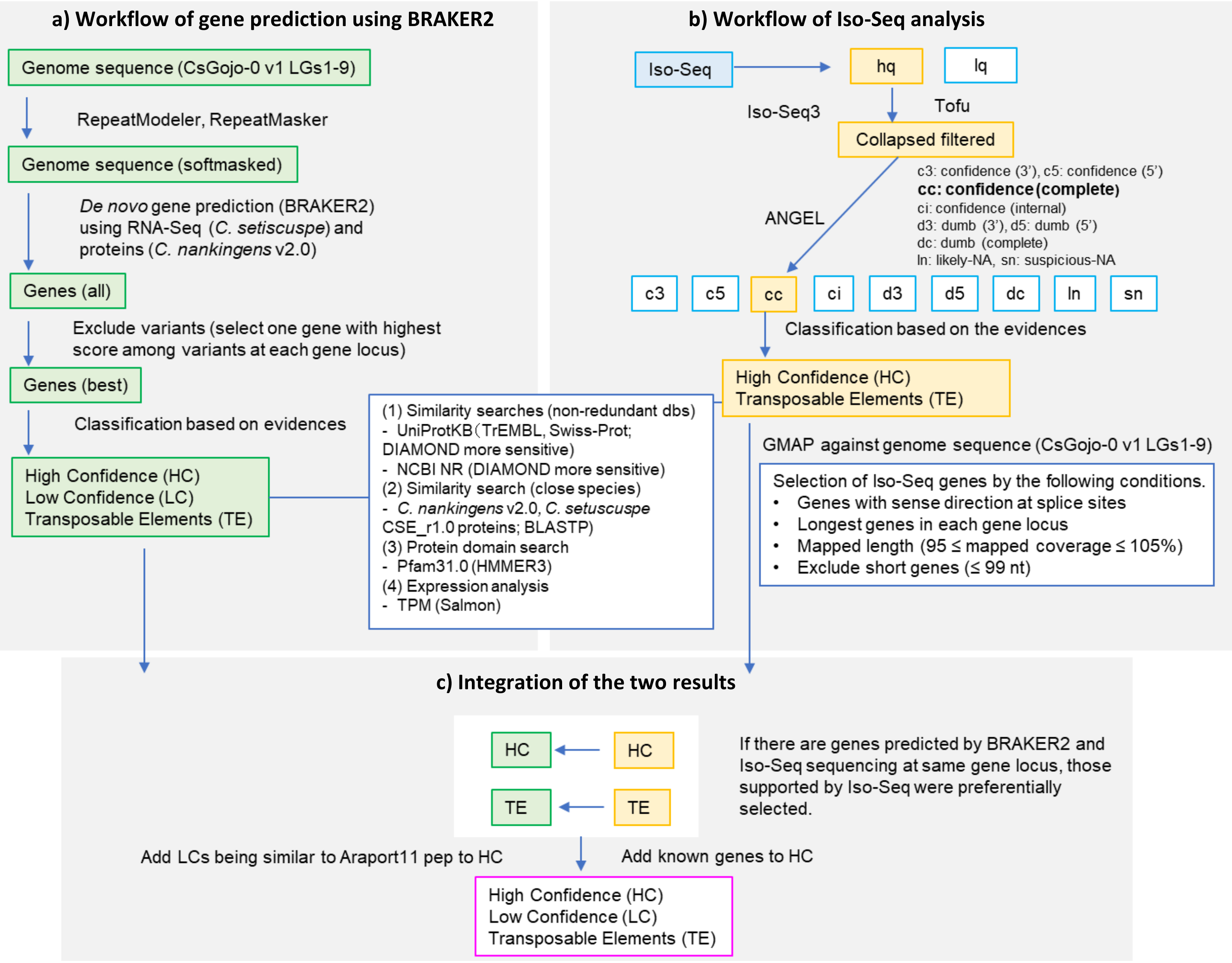


**Supplementary Figure 3 | The workflow of gene prediction. a**, The workflow of gene prediction using BRAKER2. **b**, The workflow of Iso-Seq analysis. **c**, Integration of genes predicted by BRAKER2 and Iso-Seq analysis.

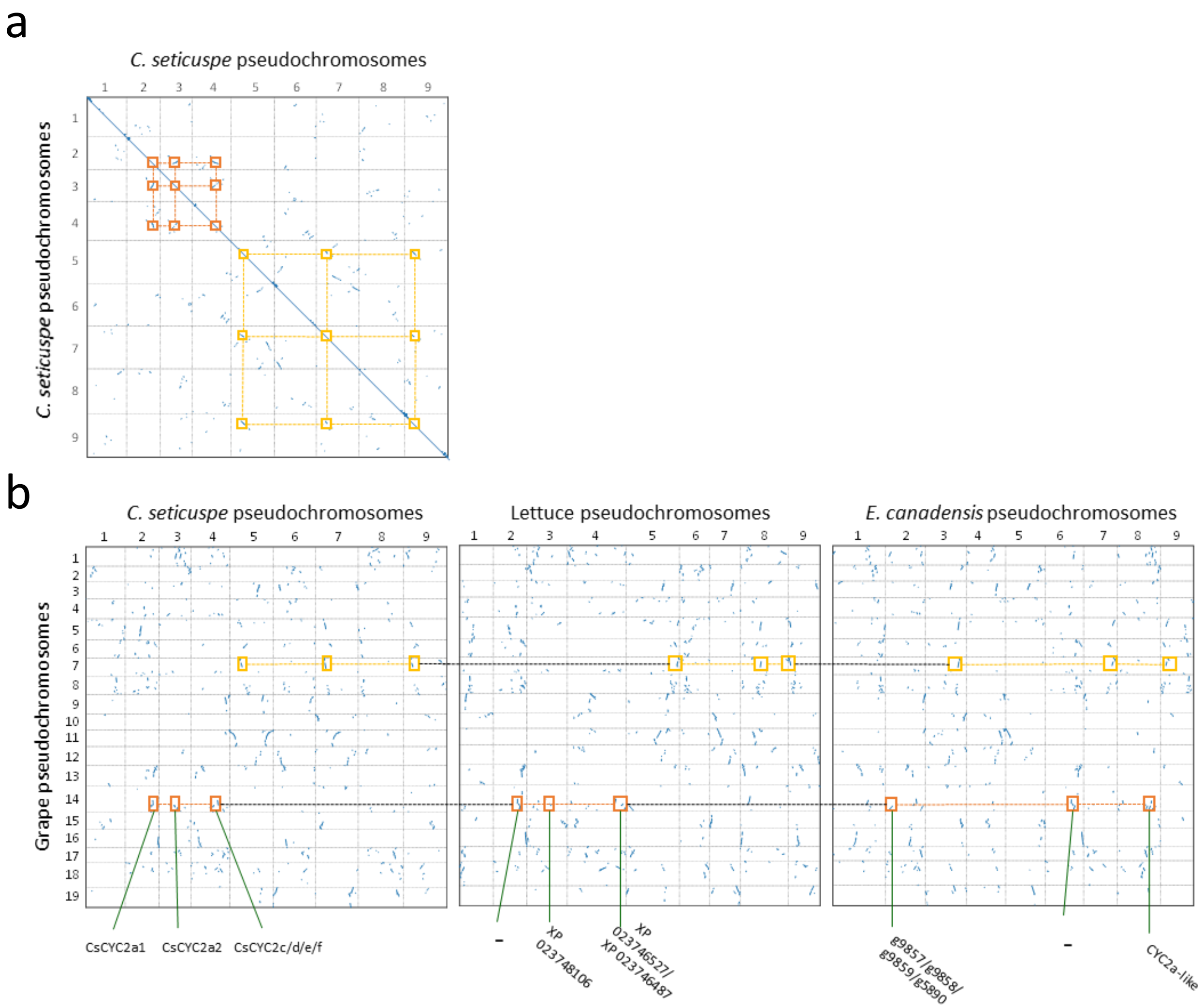


**Supplementary Figure 4 | Dot plot analysis of paralogs in *Chrysanthemum seticuspe*, lettuce, and *Erigeron canadensis*. a**, Dot plot analysis of *C. seticuspe* (Gojo-0) pseudochromosomes. The numbers represent the number of peudochromosomes/linkage groups. Some syntenic blocks between pseudochromosomes are boxed. The syntenic blocks corresponding to the same regions are represented by dotted lines. **b**, Dot plot analysis between grape (*V. vinifera)* and *C. seticuspe*, lettuce (*L. sativa*), and *E. canadensis*. The syntenic blocks corresponding to those indicated in **a** are boxed. The syntenic blocks corresponding to the same regions are represented by dotted lines. The *CYC2* family genes in the syntenic regions are shown in the dot plots.

a

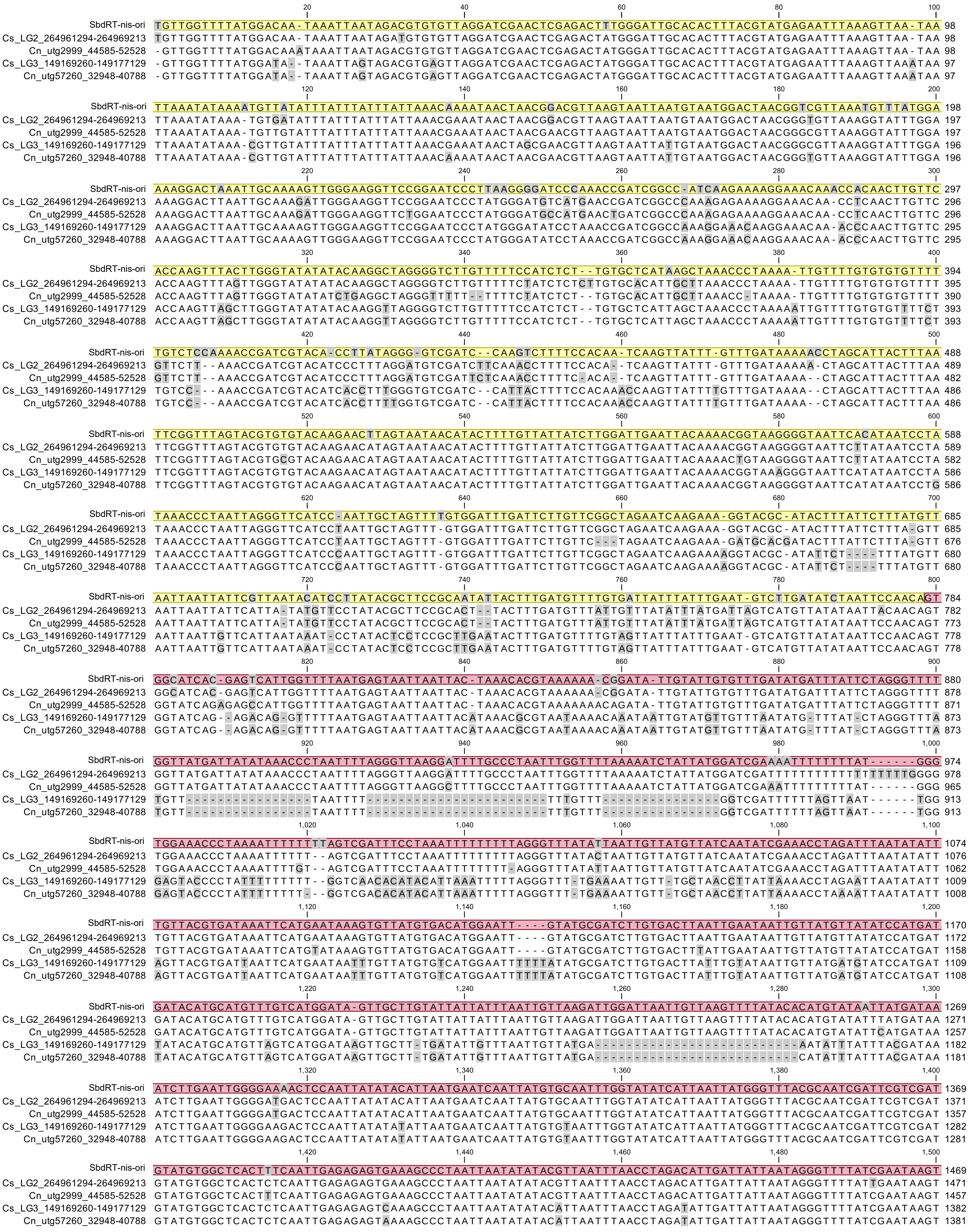
SbdRT-nis-ori
Cs_LG2_264961294-264969213
Cn_utg2999_44585-52528
Cs_LG3_149169260-149177129
Cn_utg57260_32948-40788

```
                                1,520                                  1,540                                  1,560                                  1,580                                  1,600
SbdRT-nis-ori                TGCATGTGGCTACTTGTTTACACGTATAAATTGTTTATTATGTGCGCATTGTTTTTAGTTGTTTATATTTTTTATTGGTATTACTTGTTTGTTTATTATG 1569
Cs_LG2_264961294-264969213   TGCATGTGGCTACTTGTTTACACGTATAAATTGTTTATTATGTGCGCATTGTTTTTAGTTGTTTATATTTTTTATTGGTATTACTTGTTTGTTTATTATG 1571
Cn_utg2999_44585-52528       TGCATGTGGCTACTTGTTTACACGTATAAATTGTTTATTATGTGTGCATTGTTTTTAGTTGTTTATATTTTT-ATTGGTATTACTTGTTTGTTTATTATG 1556
Cs_LG3_149169260-149177129   TGCATGTGGCTACTTGTTTACACGTATAAATTGTTTATTATGTGCGCATTGTTTTTTAGTAGTT-TATTTTTTATT--TGTTACTTGTTTGTTTATTATG 1479
Cn_utg57260_32948-40788      TGCATGTGGCTACTTGTTTACACGTATAAATTGTTTATTATA---GCATTGTTTTTTAGTAGTT-TATTTTTTATT--TGTTACTTGTTTGTTTATTATG 1475
                                1,620                                  1,640                                  1,660                                  1,680                                  1,700
SbdRT-nis-ori                TGCGACAG-TTTTAAGCAAAGGGTAATTTTGAATAGAAAAATAATACGGACTAAACAATGGCGAAATTTGATTTGTTTTGGAAGAGTTAAGTTTTCAATA 1668
Cs_LG2_264961294-264969213   TGCGACAG-TTTTAAGCAAAGGGTAATTTTGAATAGAAAAATAATACGGACTAAACAATGGCGAAATTTGATTTGTTTTGGAAGAGTTAAGTTTTCAATA 1670
Cn_utg2999_44585-52528       TGCGACAG-TTTTAAGCAAAGGGTAATTTTGAATAGAAAAATAATACGGACTAAACAATGGTGAAATTTGATTTGTTTTGGAAGAGTTAAGTTTTCAATA 1655
Cs_LG3_149169260-149177129   TTAGACAG-TTTTAAGCAAATGGTAATTTTGAATAGAAAAATAATACGGACTAAACAATGGCGAAATTTGATTTGTTTTGGAAGAGTTAAGTTTTCAATA 1578
Cn_utg57260_32948-40788      TTAGACAGGTTTTAAGCAAAGGGTAATTTTGAATAGAAAAATA-TACGGACTAAACAATGGCGAAATTTGATTTGTTTTGGAAGAGTTAAGTTTTCAATA 1574
                                1,720                                  1,740                                  1,760                                  1,780                                  1,800
SbdRT-nis-ori                GTTTAATTGATTGGTGACTTTGGTACAATGACTGAACGGCTCACCCATTCGAATGAACTATAAAGGGGTTATTTAAGAAGTGCGCACAGTCACACT-GAA 1767
Cs_LG2_264961294-264969213   GTTTAATTGATTGGTGACTTTGGTACAATGACTGAACGGCTCACCCATTCGAATGAACTATAAAGGGGTTATTTAAGAAGTGCGCACAGTCACACT-GAA 1769
Cn_utg2999_44585-52528       GTTTAATTGATTGGTGACTTTGGTACAATGACTGAACGGCTCACCCATTAGAATGAACTATAAAGGGGTTATTTAAGAAGTGC------TTACGCA-CAA 1748
Cs_LG3_149169260-149177129   GTTTAATTGATTGGTGACTTTGGTACAATGACTGAACGGCTCACCCATTCGAATGAACTATAAAGGGGTTATTTAAGAAGTGCGCACAGTCACACT-GAA 1677
Cn_utg57260_32948-40788      GTTTAATTGATTGGTGACTTTGGTACAATGG-TGAACGGCTCACCCATT-GAATGAACTATAAAGGG-TTATTTAAGAAGTGCGC-----CACACTTGAA 1666
                                1,820                                  1,840                                  1,860                                  1,880                                  1,900
SbdRT-nis-ori                AGGATGTCTAAGGTTATGCTTTTGCGCTTACCTGGTCCTCTGCCCTCTGTAAGTTCGAATGGAGCTACTTAGTAAGCCATAAAAGTACAAGCTCATCTTA 1867
Cs_LG2_264961294-264969213   TGGATGTCTAAGGTTATGCGTTTGCGCTTACCTGGTCCTCTGCCCTCTGTAAGTTCGAATGGAGCTACTTAGTAAGCCATAAAAGTACAAGCTCATCTTA 1869
Cn_utg2999_44585-52528       TGGATGTCTAAGGTTATGCGTTTGCGCTTACCTGGTCCTCTGCCCTCTGTAAGTTCGAATGGAGCTACTTAGTAAGCCATAAAAGTACAAGCTCATCTTA 1848
Cs_LG3_149169260-149177129   TGGATGTCTAAGGTTATGCGTTTGCGCTTACCTGGTCCTATGCCCTCTGTAAGTTCGAATGGAGCTACTTAGTAAGCCATAAAAGTACAAGCTCATCTTA 1777
Cn_utg57260_32948-40788      TGGATGTCTAAGGTTATGCTTTT---GCTTACACGGTCCTATGCCCTC--TAAGTTCGAATGGAGCTACTTAGTAAGCCATAAAAGTACAAGCTCATCTTA 1761
                                1,920                                  1,940                                  1,960                                  1,980                                  2,000
SbdRT-nis-ori                AGGGCGAATTAGTTTTAATTCCATGCGCCACTTCTGTGGTAATTATCCGAAACACTAAGTGTTTACACTTATGCAAAAGTACGGTATAATGGACGATAAT 1967
Cs_LG2_264961294-264969213   AGGGCGAATTAGTTTTAATTCCATGCGCCACTTCGGTGGTAATTATCCGAAACACTAAGTGTTTACACTTATGCAAAAGTACGGTATAATGGATGATGAT 1969
Cn_utg2999_44585-52528       AGGGCGAATTAGTTTTAATTCCATGCGCCACTTCGGTGGTAATTATCCGAAACACTAAGTGTTTACACTTATGCAAAAGTACGGTATAATGGATGATAAT 1948
Cs_LG3_149169260-149177129   AGGGCGAATTAGTTTTAATTCCATGCGCCACTTCGGTGGTAATTATCCGAAACACTAAGTGTTTACACTTACGCAAAAGTACGGTATAATGGATGATAAT 1877
Cn_utg57260_32948-40788      AGG-CGAATTAGTTTTAATTCCAT-CGCCACTTCGGTGGTAATTATCC-AAACACTAAGTGTTTACACTTATGCAAAAGTGCGGTATAATGGATGATAAT 1858
                                2,020                                  2,040                                  2,060                                  2,080                                  2,100
SbdRT-nis-ori                TAACACTAAATTCCTTGATTGGTGAACCCAATACCTCAATCAAAACTTTGATAATTTGATCTTTATGAATTGGTAATAAAAAGTTTTTTTATTAAAATTA 2067
Cs_LG2_264961294-264969213   TAACACTAAATTCCTTGATTGGTGAACCCAATACCTCAATCAAAACTTTGATAATTTGATCTTTATGAATTGGTAATAAAAAGTTTTTT-ATTAAAATTA 2068
Cn_utg2999_44585-52528       TAACACTAAATTCCTTAATTGGTGAACCCAATACCTCAATCAAAACTTTGATAATTTGATCTTTATGAATTGGTAATAAAAGGTTTTTT-ATTAAAATTA 2047
Cs_LG3_149169260-149177129   TAACACTAAATTCCTTGATTGGTGAACCCAATACCTCAATCAAAACTTTGATAATTTGATCTTTATGAATTGGTAATAAAAAGTTTTTT-ATTAAAATTA 1976
Cn_utg57260_32948-40788      TAACACTAAATTCCTTGATTGGTGAACCCAATACCTCAATCAAAACTTTGATAATTTGATCTTTATGAATTGGTAATAAAA-GTTTTTT-ATTAAAATTA 1956
                                2,120                                  2,140                                  2,160                                  2,180                                  2,200
SbdRT-nis-ori                TGCTACAAGATGAATGCATGACATCTTTGTACATCTTTGATTTGACCACATAAAATCAAAACTAGTACCCTTTTAAATTAAG-AATATGCCATCTAATTA 2166
Cs_LG2_264961294-264969213   TGCTACAAGATGAATGCATGACATCTTTGTACATATTTGATTTGACCACATAAAATCAAAACTAGTACCCTTTTAAATTAAGGTATATGCCATCTAATTA 2168
Cn_utg2999_44585-52528       TGCTACAAGATGAATGCATGACATCTTTGTACATCTTTGATTTGACCACATAAAATCAAAACTAATACCCTTTTAAATTAAG-TATATGCCATCTAATTA 2146
Cs_LG3_149169260-149177129   TGCTACAAGATGAATGCATGACATCTTTGTACAAATTTGATTTGACCACATAAAATCAAAACTAGTACCCTTTTAAATTAAG-AATATGCCATCTAATTA 2075
Cn_utg57260_32948-40788      TGCTACAAGATGAATGCATGACATCTTTGTACAAATTTGATTTGACCACATAAAATCAAAACTAGTACCCTTTTAAATTAAG-AATATGCCATCTAATTA 2055
                                2,220                                  2,240                                  2,260                                  2,280                                  2,300
SbdRT-nis-ori                AAAGGAACACTTGAAGTTGCAATTTACGTCATGACAGTACTTGGCGAGATTGTAATGGAGAGGTAAATCTCTATGTAAAGTCAGACCTTACATGGACATA 2266
Cs_LG2_264961294-264969213   AAAGGAACACTTGAAGTTGCAATTTAAGTCATGAGAGTACTTGGCTAGATTGTGATGGAGAGGTAAATCTCTATGTTAAGTGAGACCTTACATGGACATA 2268
Cn_utg2999_44585-52528       AAAGGAACACTTGAAGTTGCAATTTAAGTCATGAGAGTACTTGGCTAGATTGTGATGGAGAGGTAAATCTCTATGTTAAGTGAGACCTTACATGGACATA 2246
Cs_LG3_149169260-149177129   AAAGGAACACTTGAAGTTGCAATTTACGTCATGACAGTACTTGGCTAGATTGTGATGGAGAGGTAAATCTCTATGTAAAGTCAGACCTTACATGGACATA 2175
Cn_utg57260_32948-40788      AA-GGAACACTTGAAGTTGCAATTTACGTCATGAC-----TTGGCTAGATTGTGATGGAGAGGTAAATCTCTATGTAAAGTC-GACCTTACATGGACATA 2148
                                2,320                                  2,340                                  2,360                                  2,380                                  2,400
SbdRT-nis-ori                TTTGCAGTTCTTTAAGTTAGACACTTAAGTTGAGTAGGAAGTGAGA-CCTGGACTCAATGGCATAAGATGGGATGAAACATGTTTATATTTGCAATAGCG 2365
Cs_LG2_264961294-264969213   TTTGCAGTTCTTTAAGTTAGACACTTAAGTTGAGTAGGAAGTGAGATCTTGGACTCAATGGCATAAGATGGGATGAAACATGTTTATATTTGCAATAGCG 2368
Cn_utg2999_44585-52528       TTTGCAGTTCTTTAAGTTAGACACTTAAGTTGAGTAGGAAGTGAGATCTTGGACTCAATGGCATAAGATGGGATGAAACATGTTTATATTTGCAATAGCG 2346
Cs_LG3_149169260-149177129   TTTGCAGTTCTTTAAGTTAGACACTTAAATTGAGTAGGAAGTGAGA-CCTGGACTCAATGGCATAAGATGGGATGAAACATGTTTATATTTGCAATAGCG 2274
Cn_utg57260_32948-40788      TTTGCGGTTCTTTAAGTTAGACACTTAAATTGAGTAGGAAGTGAGA-CACGGACTCAATGGCATAAGATGGGATGAAACATGTTTATATTTGCAATAGCG 2247
                                2,420                                  2,440                                  2,460                                  2,480                                  2,500
SbdRT-nis-ori                TTGGAAAATATAAATCCCATAATGATCTAGGAGTTGGAAATTGTTTGCCAATTAATGTTCGACAGATTACCTTAT-TATATGCGTTCGTTGCTAGC-AGA 2463
Cs_LG2_264961294-264969213   TTGGAAAATATAAATCCCATAATGATCTAGGAATTGGAAATTGTTTGCCAATTATTGTTCGACGGATTACCCTAT-TATGTGCTTTCGTTACTAGC-AGA 2466
Cn_utg2999_44585-52528       TTGGAAAATATAAATCCCATAATGATCTAGGAATTGGAAATTGTTTGCCAATTAATGTTCGACAGATTACCTTAT-TATGTGCTTTCGTTACTAGC-AGA 2444
Cs_LG3_149169260-149177129   TTGGAAAATATAAATCCCATAATGATCTAGGAGTTGGAAATTGTTTGCCAATTAGTGTTCGACAGATTACCCTAT-TATGTGCTTTCGTTACTAGC-AGA 2372
Cn_utg57260_32948-40788      TTGGAAAATATAAATCCCATAATGATCTAGGAGTTGGAAATTGTTTGCCAATTAGTGTTCGACAGATTACCTATTATATGTGCTTTCGTTACTAGCGAGA 2347
                                2,520                                  2,540                                  2,560                                  2,580                                  2,600
SbdRT-nis-ori                TCAACTGATGTAATGAACCTATGATAATTTGGATCCTAGTTTTCGAGAAAATACTATGGTTATATGTATATTACCTTGGGTATTTTATTTTCTTGAAAAG 2563
Cs_LG2_264961294-264969213   TCAACTGATGTAATGGAACTGCGATAATTTGGATCCTAGTTTTCGAGAAAATTCTATGGTTATATGAATATTACCTTGGGTATTTTATTTTCTTGAAAAG 2566
Cn_utg2999_44585-52528       TCAACTGATGTAATGGAACTATGATAATTTGGATCCTAGTTTTCGAGAAAATTCTATGGTTATATGAATATTACCTTGGGTATTTTATTTTCTTGAAAAG 2544
Cs_LG3_149169260-149177129   TCAACTGATGTAATGGAACTACGATAATTTGGATCCTAGTTTTCGAGAAAATTCTATGGTTATATGAATATTACCTTGGGTATTTTATTTTCATGAAAAG 2472
Cn_utg57260_32948-40788      TCAAC-GATGTAATGGAACTATGATAATTTGGATCCTAGTTTTCGAGAAAATTCTATGGTTATATGAATATTACCTTGGGTATTTTATTTTCTTGAAAAG 2446
                                2,620                                  2,640                                  2,660                                  2,680                                  2,700
SbdRT-nis-ori                TTTAATTATCGAAAATACTAAAATTGAATTTTGCTAATTCTTTAAATGTATTAATGCAACATTAAACCGTAAACGAAGGTTATTCCTGTATTGGTAAA-T 2662
Cs_LG2_264961294-264969213   TTTAATTATCGAAAATACTAAAATTGAATTTTGCTAATTCTTTAAATGTATTAATGCAACATTAAACCGTAAACGTAGGTTATTCATGTATTGGTAAAAT 2666
Cn_utg2999_44585-52528       TTTAATTATCGAAAATACTAAAATTGAATTTTGCTAATTCTTTAAATGTATTAATGCAACATTAAATCGTAAACGTAGGTTATTCTTGTATTGGTAAA-T 2643
Cs_LG3_149169260-149177129   TTTAATTATCGAAAATACTAAAATTGAATTTTGCTAATTCTTTAAATGTATTAATGCAACATTAAATCGTAAACGTAGGTTATTCTTGTATTGGTAAA-T 2571
Cn_utg57260_32948-40788      TTTAATTATCGAAAATACTAAAATTGAATTTTGCTAATTCTTTAAATGTATTAATGCAACATTAAATCGTAAACGTAGGTTATTCTTGTATTGGTAAA-T 2545
                                2,720                                  2,740                                  2,760                                  2,780                                  2,800
SbdRT-nis-ori                TGTTCTACACCTGTAAGGGTTTT-TGGTAGAGTCAACTTTACTAAAAGAATAAACCTAGTCCTCTTTTC-TAAGAAAGAGTGAGATTCAATCCTATTCCT 2760
Cs_LG2_264961294-264969213   TGATCTACACCTGTAAGGGTTT--TGGTAGAGTCAACTTTACTAAAAGAATAAACCTAGTCCTCTTTTC-TAAGAAAGAGTGAGATTCAACCCTATTCCT 2763
Cn_utg2999_44585-52528       TGATCTACACCTGTAAGGGTTTTTTGGTAGAGTCAACTTTACTAAGAGAATAAACCTAGTCCTCTTTTCATAAGAAAGAGTGAGATTCAACCCTATTCCT 2743
Cs_LG3_149169260-149177129   TGATCTACACCTGTAAGGGTTTT-TGGTAGAGTCAAATTTACTAAAAGAATAAACCTAGTCCTCTTTTCACGCGAAAGAGTGAGATTCAACCCTATTCCT 2670
Cn_utg57260_32948-40788      TGATCTACACCTGTAAGGGTTTT-TGGTAGAGTCAACTTTACTAAAAGAATAAACCTAGTCCTCTTTTCACGCGAAAGAGTGAGATTCAACCCTATTCCT 2644
                                2,820                                  2,840                                  2,860                                  2,880                                  2,900
SbdRT-nis-ori                TTTCATGAATGAAGAAAGGTATGATAGGACAATTAGAATTGTCTAAATTCTAAATACATAACTCGTAAGCAAATTTTAATAGTTAAGAAAATTGCATATC 2860
Cs_LG2_264961294-264969213   TTTCATGAATGAAGAAAGGTATGATAGGACAATTAGAATTGTCTAAATTCTAAATACATAACTCGTAAGCAAATTTTAATAGTTAAGAAAATTGCATATT 2863
Cn_utg2999_44585-52528       TTTCATGAATGAAGAAAGGTATGATAGGACAATTAGAATTGTCTAAATTCTAAATACATAACTCGTAAGCAAATTTTAATAGTTAAGAAAATTGCATATT 2843
Cs_LG3_149169260-149177129   TTTCATGAATGAAGAAAGGTATGATAGGACAATTAGAATTGTCTAAATTCTAGATACATAACTCGTAAGCGAATTTTAATAGTTAAGAAAATTGCATATT 2770
Cn_utg57260_32948-40788      TTTCATGAATGAAGAAAGGTATGATAGGACAATTAGAATTGTCTAAATTCTAGATACATAACTCGTAAGCGAATTTTAATAGTTAAGAAAATTGCATATT 2744
                                2,920                                  2,940                                  2,960                                  2,980                                  3,000
SbdRT-nis-ori                TCTGAGTTATTAAATTAGAAGCTAGACTAATTACTGTATAATTGGTAAAAGACAAACTAGCATATTTTTATGTAGTTGTT-TGTTTTGACCATTTAAAAGT 2959
Cs_LG2_264961294-264969213   TCTGAGTTATTAAATTAGAAGCTAGACTAATTACTGTATAATTGGTAAAAGACAAACTAACATATTTTTATCTAGTTGTTGTGCTTTGACCATTTAAAAGT 2963
Cn_utg2999_44585-52528       TCTGAGTTATTAAATTAGAAGCTAGACTAATTACTGTATAATTGGTAAAAGACAAACTAGCATATTTTTATCTAGTTGTTTTGTTTTGACCATTTAAAAGT 2943
Cs_LG3_149169260-149177129   TCTGAGTTATTAAATTAGAAGCTAGACTAATTACTGTATAATTGGTAAAAGACAAACTAGCATATTTTTATCTAGTTGTTGCACTTTGACCATTTAAAAGT 2870
Cn_utg57260_32948-40788      TTTGAGTTATTAAATTAGAAGCTAGACTAATTACTGTATAATTGGTAAAAGACAAACTAGCATATTTTTATCTAGTTGTTGTACTTTGACCATTTAAAAGT 2844
```

```
                              3,020     3,040     3,060     3,080     3,100
SbdRT-nis-ori                 ACTACTTACAATTGTTGGAATAGTAGAACTATTAAGAAAGAAATTGCATGCCCAATTTTTCAAGTTAATAGCATTGATGAATAA-TTCATCATCTACATA 3058
Cs_LG2_264961294-264969213    ACTACTTACAATTGTTGGAATAGTAGAACTATTAAGAAAGAAATTGCATGCCCAATTTTTCAAGTTAATAGCATTGATGGATAG-TTCATCATCTACATA 3062
Cn_utg2999_44585-52528        ACTACTTACAATCGTTGGAATAGTAGAACTATTAAGAAAGAAATTGCATGCCCAAATTTTCAAGTTAATAGCATTGATGGATAAATTCATCATCTACATA 3043
Cs_LG3_149169260-149177129    ACTACTTACAATTGTTGGAATAGTATAACTATTAAGAAAGAAATTACATGCCCAAATTTTCAAGTTAATAGCATTGATGGATAAATTCATCATCTACATA 2970
Cn_utg57260_32948-40788       ACTACTTACAATTGTTGGAATAGTAGAACTATTAAGAAAGAAATTACATGCCCAATTTTTCAAGTTAATAGCATTGATGGATAAATTCATCATCTACATA 2944

                              3,120     3,140     3,160     3,180     3,200
SbdRT-nis-ori                 GTAGAATACATTTTTTTAAGTACATAAGTAATTGTTATGTATGGACTTTCTCAAAGGTGTAGG-AAGTAAATAATACACTATTTTCTTTCACTGGTTAAT 3157
Cs_LG2_264961294-264969213    GTAGAATACACTCTTT-AAGTACATAAGTAATTGTTATGTATGGACTTTTTCAAAGGTGTAGGGAAGTAAATAGAACACTATTTTCTTTTACTGGTTAAT 3161
Cn_utg2999_44585-52528        GTAGAATACATTCTTTTAAGTACATAAGTAATTGTTATGTATGGACTTTCTCAAAGGTGTAGGGAAGTAAATAGAACACTATTTTCTTTTACTGGTTAAT 3143
Cs_LG3_149169260-149177129    GTAGAATACATTCTTTTAAGTACATTTGTAATTGTTATGTATGGACTTTCTCAAAGGTGTAGG-AAGTAAATAATACACTATTTTCTTTCACTGGTTAAT 3069
Cn_utg57260_32948-40788       GTAGAATACATTCTTTTAAGTACATTTGTAATTGTTATGTATGTACTTTCTCAAAGGTGTAGG-AAGTAAATAATACACTATTTTCTTTCACTGGTTAAT 3043

                              3,220     3,240     3,260     3,280     3,300
SbdRT-nis-ori                 GAAACACTTACAATGTTGTCAAATCCAGTGGGAGTGTTGTGTTCTTATTACTGAATAAGGAATTAATGTAATGATCTCTTATTCAGTGGGAGTTACAAGC 3257
Cs_LG2_264961294-264969213    GAAACACTTCCAATGTTGTCAAATCCAGTGGGAGTATGTTGTTCTTATTACTGAATAAGGACTTAATGTAATGATCTCTTATTCAGTGGGAGTTACAAGC 3261
Cn_utg2999_44585-52528        GCAACACTTACAATGTTGTCAAATCCAGTGGGAGTGTTGTGTTCTTATTACTGAATAAGGAATTAATGTAATGATCTCTTATTCAGTGGGAGTTACAAGC 3243
Cs_LG3_149169260-149177129    GAAACACTTACAATGTTGTCAAATCCAGTGGGAGTGTTGTGTTCTTATTACTGAATAAGGAATTAATGTAATGATCTCTTATTTAGTGGGAGTTACAAGC 3169
Cn_utg57260_32948-40788       GAAACACTTACAATGTTGTCAAATCCAGTGGGAGTGTTGTGTTCTTATTACTGAATAAGGAATTAATGTAATTATCTCTTATTCAGTGGGAGTTACAAGC 3143

                              3,320     3,340     3,360     3,380     3,400
SbdRT-nis-ori                 TTTGTCATATTGGTAGTAACCCAGATTATCTCTAGAATGTCTTAAGATAATTACTTTCTACTTGTAGACACTTATGTTTTCAAGGATAGTCATGTACGTT 3357
Cs_LG2_264961294-264969213    TTTGTCATATTGGTAGTAACTCAGATTATCTCTAGAATGTCTTATGATAATTACTTTCCACTTGTAGACACTTATGTTTTCAAGGATAGTCATGTACGTT 3361
Cn_utg2999_44585-52528        TTTGTCATCTTGGTAGTAACCCAGATTATCTCTAGAATGTCTTAAGATAATTACTTTCCACTTGAAGACACTTATGTTTTCAAGGATAGTCATGTACGTT 3343
Cs_LG3_149169260-149177129    TTTGTCATCTTGGTAGTAACCCATATTATCTCTTGAATGTCTTAAGATAATTACTTTCCACTTATAGACACTTATGTTTTCAAGGATAGTCATGTACGTT 3269
Cn_utg57260_32948-40788       TTTGTCATCTTGGTAGTAACCCAGATTATCTCTAGAATGTCTTAAGATAATTACTTTCCACTTATAGACACTTATGTTTTCAAGGATAGTCATGCACGTT 3243

                              3,420     3,440     3,460     3,480     3,500
SbdRT-nis-ori                 TCCTTATTAGACAATAAAGTGGTTCTACATCTACTAGTTATGCTAATATCAATGACACAATCATATTAGTCACATAAACAAACATATGTCGTGTTGTTTA 3457
Cs_LG2_264961294-264969213    TCCTTATTAGACAATGAAGTGATTCTACATCTACTAGTTATGCTAATATCAATGACACAATCATATTAGTCACATAAACAAACATATGTCGTGTTGTTTA 3461
Cn_utg2999_44585-52528        TCCTTATTAGACAATGAAGTGGTTCTACATCTACTAGTTATGCTAATATCAATGACACAATCATATTAGTCACATAAACAAACATATGTCGTGTTGTTTA 3443
Cs_LG3_149169260-149177129    TCCTTATTAGACAATGAAGTGGTTCTACATCTACTAGTTATGCTAATATCAATGACACAATCATATTAGTCACATAAACAAACATATGTCGTGTTGTTTA 3369
Cn_utg57260_32948-40788       TCCTTATTAGACAATGAAGTGGTTCTACATCTACTAGTTATGCTAATATCAATGACACAATCATATTAGTCACATAAACAAACATATGTCGTGTTGTTTA 3343

                              3,520     3,540     3,560     3,580     3,600
SbdRT-nis-ori                 TTTGCTACATGATGTTGTTCCATGCACAAGTGGGAGAAATCTATATTGTGTGCAAGGTACAACATATAGTTATATGATTTAATAGATTCTCTTGACTGCA 3557
Cs_LG2_264961294-264969213    TCTGCTACATGATGTTGTTCCATGCACAAGTGGGAGAAATTTATATTGTGTGCAAGGTACAACATATAAGTATATGATTTAAAGGATTCTCTTGACTGCA 3561
Cn_utg2999_44585-52528        TCTGCTACATGATGTTGTTACATGCACAAGTGGGAGAAATCTATATTGTGTGCAAGGTACAACATATAAGTATATGATTTAACGGATTCTCTTGACTGCA 3543
Cs_LG3_149169260-149177129    TCTGCTACATGATTTTGTTTCATGCACAAGTGGGAGAAATCTATATTGTGTGCAAGGGACAACATATAGGTATATGATTTAAAGGATTCTCTTGACTGCA 3469
Cn_utg57260_32948-40788       TCTGCTACATGATTTTGTTCCATGCACAAGTGGGAGAAATCTATATTGTGTGCAAGGGACAACATATAGGTATATGATTTAAAGGATTCTCTTGACTGCA 3443

                              3,620     3,640     3,660     3,680     3,700
SbdRT-nis-ori                 AGGTAAGAGACTAACATTCTAATATAACTTGTACATATCTAATTCAGGAAACATTTGGTGTTGCAATTTACGTCATGTCAGAACTTGGCGAGATTGCAAT 3657
Cs_LG2_264961294-264969213    AGGTAAGAGACTAACGTTCTAATATAACTTGTACATATCTAATTCAAGAAACACTTGGTGTTACAATTTACGTCATGTGAGAACTTGGCGAGATTGCAAC 3661
Cn_utg2999_44585-52528        AGGTAAGAGACTAACGTTCTAATATAACTTGTACATATCTAATTCAAGAAACACTTGGTGTTACAATTTACGTCATGTGAGAACTTGGCGAGATTGCAAC 3643
Cs_LG3_149169260-149177129    AGGTAAGAAACTAACGTTCTAATATAAATTGTACATATCTAATTCAAGAAACACTTGGTGTTGCAATTTACGTCATGTCAGAACTTGGCGAGATTGCAGC 3569
Cn_utg57260_32948-40788       AGGTAAGAAACTAACGTTCTAATATAACTTGTACATATCTAATTCAAGAACCACTTGGTGTTGCAATTTACGTCATGTCAGAACTTGGCGAGATTGCAGC 3543

                              3,720     3,740     3,760     3,780     3,800
SbdRT-nis-ori                 AACAAAGTAAATCTCTATGTGAAGTCAGACCTTGCATGGACATATTTACAGTTTCTAAATTAGACAAGTATTTAGAACATGTGTCAAAATGATATATAAG 3757
Cs_LG2_264961294-264969213    AACAAAGTAAATCTCTATGTGAAGTCAGACCTTGCATGGACATATTTGCAGTTTCTAAATTAGACAAGTATTTAGAACATGTGTCAAAATGATATATAAG 3761
Cn_utg2999_44585-52528        AACAAAGTAAATCTCTATGTAAAGTGAGATCTTGCATGGACATATTTACAGTTTCTAAATTAGACAAGTATTTAGAACATGTGTCAAAATGATATATAAG 3743
Cs_LG3_149169260-149177129    AACAAAGTAAGTCTCTATGTGAAGTCAGACCTTGCATGGACATATTTGCAGTTTCTAAATTAGACAAGTATTTAGAACATGTGTCAAAATGATATATAAG 3669
Cn_utg57260_32948-40788       AACAAAGTAAGTCTCTATGTGAAGTCAGACCTTGCATGGACATATTTGCAGTTTCTAAATTAGACAAGTATTTAGAACATGTGTCAAAATGATATATAAG 3643

                              3,820     3,840     3,860     3,880     3,900
SbdRT-nis-ori                 ACTACTTCAAATTAAATATTTATGTATTATTAGAATGACATAAAATTAAATAAAGTAATATAATCCTACTTCAA-TTTGTTTATTTATGTATTTTGGAAT 3856
Cs_LG2_264961294-264969213    ACTACTTCAAATTAGATTTTTATGTATTATTAGAATGACATAAA------TGAAGTAATCTAATACTACTTCAAATTTGTTTATTTATGTATTTTGGAAT 3855
Cn_utg2999_44585-52528        ACTACTTCAAATTAGATTTTTATGTATTATTAGAATGACATAAA------TGAAGTAATCTAATACTATTTCAAATTTGTTTATTTATGTATTTTGGAAT 3837
Cs_LG3_149169260-149177129    ACTACTTCAAATTAAATATTTATGTATTATCAGAATGACATAAAATTAAATAAAGTAATCTAATCCTACTTCAA-TTTGTTTATTTATGTATTTTGGAAT 3768
Cn_utg57260_32948-40788       ACTACTTCAAATTAAATATTTATGTATTATTAGAATGACATAAAATTAAATAAAGTAATCTAATCCTACTTCAA-TTTGTTTATTTATGTATTTTGGAAT 3742

                              3,920     3,940     3,960     3,980     4,000
SbdRT-nis-ori                 GACATAAATGAAAAATAATGTAGTCCTACATTATTATGAAGTAAAATAAATTAACTTGAAATTACTATTGAAACCAAGAGTTGTGTTCATAATTACTTTA 3956
Cs_LG2_264961294-264969213    GACATAAATGAAAAATAATGTAGTCCTGCATTATTGTGAAGTAAAATAAATTAACTTGAAATTACTATTGAAACCAAGAGTTGTGTTCATAATTACTTTA 3955
Cn_utg2999_44585-52528        GACATAAATGAAAAATAATGTAGTCCTACATTATTGTGAAGTAAAATAAATTAACTTGAAATTACTATTGAAACCAAGAGTTGTGTTCATAATTACTTTA 3937
Cs_LG3_149169260-149177129    GACATAAATGAAAAATAATGTAGTCCTACATTATTATGAAGTAAAATAAATTAACTTGAAATTACTATTGAAACCAAGAGTTGTGTTCATAATTACTTTA 3868
Cn_utg57260_32948-40788       GACATAAATGAAAAATAATGTAGTCCTACATTATTATGAAGTAAAATAAATTAACTTGAAATTACTATTGAAACCAAGAGTTGTGTTCATAATTACTTTA 3842

                              4,020     4,040     4,060     4,080     4,100
SbdRT-nis-ori                 GTTAAACATGTTTATGATTCATTGTAAAGACATAAACATAAAGTCTTTTAAGACTTATAATAAAGTAAGACGTTAACTTGTTATGTTTTTT-AATTAAGA 4055
Cs_LG2_264961294-264969213    GTTAAACATGTTTATGATTCATTGTAAAGACATAAACATAAAGTCTTTTAAGACTTATAATAAAGTAAGACGTTAACTTGTTATGTTTTTT-AATTAAGA 4054
Cn_utg2999_44585-52528        GTTAAACATGTTTATGATTCATTGTAAAGACATAAACATAAAGTCTTT-AAGACTTATAATAAAGTAAGACGTTAACTTGTTATGTTTTTT-AATTAAGA 4035
Cs_LG3_149169260-149177129    GTTAAACATGTTTATGATTCATTGTAAATACATAAACATAAAGTCTTTTAAGACTTATAATAAAGTAAGACGTTAACTTGTTATGTTTTTT-AATTAAGA 3967
Cn_utg57260_32948-40788       GTTAAACATGTTTATGATTCATTGTAAAGACATAAACATAAAGTCTTTTAAGACTTATAATAAAGTAAGACGTTAACTTGTTATGTTTTTTTAATTAAGA 3942

                              4,120     4,140     4,160     4,180     4,200
SbdRT-nis-ori                 CATATACAAGAAGTCATTGAGTTGATCAATGATTTATAACGTATAATGGGATTATAACAGTTTATGTTTCTGGTTGAGACATGGATATGAAGTGACTTGG 4155
Cs_LG2_264961294-264969213    CATATACAAGAAGTCAT---------CAATGATTTATAACGTATAATGGGATTATAACAGTTTATGTTTCTTGTTGAGACATGGATATGAAGTGACTTGG 4145
Cn_utg2999_44585-52528        CATATACAAGAAGTCATTGAGTTGATCAATGATTTATAACGTATAATGGGATTATAACAGTTTATGTTTCTTGTTGAGACATGGATATGAAGTAACTTGG 4135
Cs_LG3_149169260-149177129    CATATACAAGAAGTCATTGAGTTGATCAATGATTTATAACGTATAATGGGATTATAACAGTTTATGTTTCTTTTTGAGACATGGATATGAAGTGACTTGG 4067
Cn_utg57260_32948-40788       CATATACAAGAAGTCATTGAGTTGATCAATGATTTATAACGTATAATGGGATTATAACAGTTTATGTTTCTTGTTGAGACATGGATATGAAGTGACTTGG 4042

                              4,220     4,240     4,260     4,280     4,300
SbdRT-nis-ori                 TTAAAAGGTACTTGTGTTATAAAGCATGCAATTGGCTTGGAAAGTCTATTATACCATTAAGTTGACTTAGTTGGTAAATAGGTAAGCCAAAAGGAATATG 4255
Cs_LG2_264961294-264969213    TTAAAAGGTACTTGTGTTATAAAGCATGCAATTGGCTTGGAAAGTCTATTATACCATTAAGTTGACTTAGTTGGTAAATAGGAAATTCAAAATGAATATG 4245
Cn_utg2999_44585-52528        TTAAAAGGTACTTGTGTTATAAAGCATGCAATTGGCTTGGAAAGTCTATTATACCATTAAGTTGACTTAGTTGGTAAATAGGTAATTCAAAATGAATATG 4235
Cs_LG3_149169260-149177129    TTAAAAGGTACTTGTGTTATAAAGCATGTAATTGGCTTGAAAAGTCTATTATACCATTAAGTTGACTTAGTTGGTAAATAGGTAAGCCAAAAGGAATATG 4167
Cn_utg57260_32948-40788       TTAAAAGGTACTTGTGTTATAAAGCATGTAATTGGCTTGAAAAGTCTATTATACCATTAAGTTGACTTAGTTGGTAAATAGGTAAGCCAAAAGGAATATG 4142

                              4,320     4,340     4,360     4,380     4,400
SbdRT-nis-ori                 TTTATCTTATATATTGTTAGATTGTCTTACAATATTAAA-TTCTAATGAATACCACAAGTTGATGGTATGTTCAGTTAAGAGAATCAAACTCCATTTGAT 4354
Cs_LG2_264961294-264969213    TTTATCTTATATATTGTTAGATTGTCTTACAATATTAAAATTCTAATGAATACCACAAGTTGATGGTATGTTCAGTTAGGAGAATCAAACTCTATTTGAT 4345
Cn_utg2999_44585-52528        TTTATCTTATATATTGTTAGATTGTCTTACAATATTAAA-TTCTAATGAATACCACAAGTTGATGGTATGGTCAGTTAGAAGAATCAAACTCTATTTGAT 4334
Cs_LG3_149169260-149177129    TTTATCTTATATATTGTTAGATTGTCTTACAATATTAAA-TTCTAATGAATACCACAAGTTGATGGTATGTTCAGTTAGGAGAATCAAACTCTATTTGAT 4266
Cn_utg57260_32948-40788       TTTATCTTATATATTGTTAGATTGTCTTACAATATTAAA-TTCTAATGAATACCACAAGTTGATGGTATGTTCAGTTAGGAGAATCAAACTCTATTTGAT 4241

                              4,420     4,440     4,460     4,480     4,500
SbdRT-nis-ori                 TTGTTTGATTCATGATAACCATAACTTCTTTCTTAAAAAGTAACAAGGTTAGAATATTGAGTATGGATTGAATATACTCAATATGACTCCAACCTAGAAG 4454
Cs_LG2_264961294-264969213    TTGTTTGATTCAAGATAACCATAACTTCTTTCTTAAAAAGTAACAAGGTTAGAATATTGAGTATGGATTGAATATACTCAATATGACTCCAACCTAGAAG 4445
Cn_utg2999_44585-52528        TTGTTTGATTCAAGATAACCATAACTTCTTTCTTAAAAAGTAACAAGGTTAGAAAATTGAGTATGGATTGAATATACTCAATATGACTCCAACCTAGAAG 4434
Cs_LG3_149169260-149177129    -----TGATTCAAGATAACCATAACTTCTTTCTTAAAAAGTAACAAGGTTAGAAAATTGAGTATGGATTAAATATACTCAATATGACTCCAACCTAGAAG 4361
Cn_utg57260_32948-40788       -----TGATTCAAGATAACCATAACTTCTTTCTTAAAAAGTAACAAGGTTAGAAAATTGAGTATGGATTAAATATACTCAATATGACTCCAACCTAGAAG 4336
```

```
                                     4,520               4,540               4,560               4,580               4,600
SbdRT-nis-ori                 ATTAATAAAACA-CCATTTCATTATGGAATGTGATGAATCACATTAAGTTTTAAAGTTGAAGATTCTGAATTTGTTAAACTACTTATGAATAGGTTATTA 4553
Cs_LG2_264961294-264969213    ATTAATAAAACA-CCATTTCATTATGGAATGTGATGAATCACATTAAGTTTTAAAATTGAAGATTCTGAATTTGTTAAACTACTTATGAATAGGTTATTA 4544
Cn_utg2999_44585-52528        ATTAATAAAACA-CCATTTCATTATGGAATGTGATGAATCACATTAAGTTTTAAAATTGAAGATTCTGAATTTGTTAAACTACTTATGAATAGGTTATTA 4533
Cs_LG3_149169260-149177129    ATTGATAAAAAAACCATTTTATTATGGAATGTGATGAATCACATTAAGTTTTAAAATTGAAGATTCTGAATTTGTTAAACTACTTATGAATAGGTTATTA 4461
Cn_utg57260_32948-40788       ATTGATAAAACA-CCATTTTATTATGGAATGTGATGAATCACATTAAGTTTTAAAATTGAAGATTCTGAATTTGTTAAACTACTTATGAATAGGTTATTA 4435
                                     4,620               4,640               4,660               4,680               4,700
SbdRT-nis-ori                 ACAAACTAATGTCAAGAATTATCAATATGTATTTGTAATGTCCCAAATAGAGAACAATGTAGTTTTGCTCTAATTGCTAAGTATATATAGATAGTCTATA 4653
Cs_LG2_264961294-264969213    ACAAACTAATGTCAAGAATTATCAATGTGTATTTGTAATGTCCCAAGTAGAGAACAATGTAGTTTTGCTCTAATTGCTAAGTATATATAGATAGTCTATA 4644
Cn_utg2999_44585-52528        ACAAACTTAAGTCAGGAATTATCAATGTGTATTTGTAATGTCCCAAATAGAGAACAATGTAGTTTTGCTCTAATTGCTAAGTATATATAGATAGTCTATA 4633
Cs_LG3_149169260-149177129    ACAACGTAAAGTCAGGAATTATCAATGTGTATTTGTAATGTCCCAAAAAGAGAACAATGTAGTTTTGCTCTAATTGCTAAGTATATATAGATAGTCTATA 4561
Cn_utg57260_32948-40788       ACAACGTAAAGTTAGGAATTATCAATGTGTATTTGTAATGTCCCAAAAAGAGAACAATGTAGTTTTGCTCTAATTGCTAAGTATATATAGATAGTCTATA 4535
                                     4,720               4,740               4,760               4,780               4,800
SbdRT-nis-ori                 TAT--TACAAAAGGCAAGTGGGAGTTTTTTTATCTTGAAAGATAAAATTACAAGATACATAATCTTCTAAAACACACTAGTAAATATCATGATGAGGTTA 4751
Cs_LG2_264961294-264969213    TAT--TACAAAAGGCAAGTGGGAGTTTTTT-ATCTTGAAAGATAAAATTACAAGATACATAATCTTCTAAAACACACTAGTAAATATCATGATGAGGTTA 4741
Cn_utg2999_44585-52528        TAT--TACAAAAGGCAAGTGGGAGTTTTTT-ATCTTGAAAGATAAAATTACAAGATACATAATCTTCTAAAACACACTAGTAAATATCATGATGAGGTTA 4730
Cs_LG3_149169260-149177129    TATATTACAAAAGGCAAGTGGGAGTTTT---ATCTTGAAAGATAAAATTACAAGATACATAATCTTCTAAAACACACTAGTAAATATCATGATGAGGTTA 4658
Cn_utg57260_32948-40788       TATATTACAAAAGGCAAGTGGGAGTTTT---ATCTTGAAAGATAAAATTACAAGATACATAATCTTCTAAAACACACTAGTAAATATCATGATGAGGTTA 4632
                                     4,820               4,840               4,860               4,880               4,900
SbdRT-nis-ori                 GACATGTAATTGATAGACCTCATGATGATGGAATTTTCATTCATAGATCTGCAAGGATATCTCAGATATTTGACTTATATGCTGAGAAATATGAGTTACA 4851
Cs_LG2_264961294-264969213    GACATGTAAATGATAGACCTCATGATGATGGAATTTTCATTCATAGATCTGCAAAGATATCTCAGATATTTGACTTATATGCTGAGAAATATGAGTTACA 4841
Cn_utg2999_44585-52528        GACATGTAAATGATAGACCTCATGATGATGGAATTTTCATTCATAGATCTGCAAGGATATCTCATATATTTGATTTATATGCTGAGAAATATGAGTTACA 4830
Cs_LG3_149169260-149177129    GACATGTAAATGATAGACCTCATGATGATGGAATTTTCATTCATAAATCTGCAAGGATATCTCAGATATTTGACTTATATGCTGAGAAATATGAGTTACA 4758
Cn_utg57260_32948-40788       GACATGTAAATGATAGACCTCATGATGATGGAATTTTTATTCATAGATCTGCAAGGATATCTCAGATATTTGACTTATATGCTGAGAAATATGAGTTACA 4732
                                     4,920               4,940               4,960               4,980               5,000
SbdRT-nis-ori                 GGATTATAGTAAACCTGATAATCATTATGATGGATTACCAAATATTGAATTGGGTAAATTACTCATACTATGAATGTTAAAGATGCTGTCCATAAAGATA 4951
Cs_LG2_264961294-264969213    GGATTATAGTAAACCTGATAATCATTATGATGGATTACCAAATATTGAATTGGGAAAATTACTCATACTCTGAATGTTAAAGATGCTGTCCATAAAGATA 4941
Cn_utg2999_44585-52528        GGATTATAGTAAACCTGATAATCATTATGATGGATTACCAAATTTTGAATTAGGAAAATTACTCATACTATGAATGTTAAAGATGCTGTCCATAAAGATA 4930
Cs_LG3_149169260-149177129    GGATTATAGTAAACCTGATAATCATTATGATGGATTACCGAATGTTGAATTGGGTAAATTACTCATACTATGAATGTTAAAGATGCTGTCCATAAAGATA 4858
Cn_utg57260_32948-40788       GGATTATAGTAAACCTGATAATCATTATGATGGATTACCGAATGTTGAATTGGGTATATTACTCATACTATGAATGTTAAAGATGCTGTCCATAAAGATA 4832
                                     5,020               5,040               5,060               5,080               5,100
SbdRT-nis-ori                 ATCAATTTTGGAAATTTGGTTTATCTTAATAGTTAGACTAGAGAATGGAAATGAGCTTCTTCAAA-GAGATGAAAAACCCTAGTGGTTTGAATTTCTATG 5050
Cs_LG2_264961294-264969213    ATCAATTTTGGAAATTTGATTTATCTTAATAG---------AGAATGGAAATGAGTTTCTTCAAAAGAGATGAAAAACCCTTGTGGTTTGAATTTCTATG 5032
Cn_utg2999_44585-52528        ATCAATTTTTGAAATTTGGTTTATCTTAATAGTTAGACTAGAGAATGGAAATGAGTTTCTTCAAA-GAGATGAAAAACCCTAGTGGTTTGAATTTCTATG 5029
Cs_LG3_149169260-149177129    ATCAATTTTGGAAATTTGGTTTATCTTAATAGTTAGACTAGAGAATGGAAATGAGTTTCTTCAAA-GAGATGAAAAACCCTAGTGGTTTGAATTTCTATG 4957
Cn_utg57260_32948-40788       ATCAATTTTGGAAATTTGGTT-ATCTTAATAGTTAGACTAGAGAATGGAAATGAGTTTCTTCAAA-GAGATGAAAAACCCTAGTGGTTTGAATTTCTATG 4930
                                     5,120               5,140               5,160               5,180               5,200
SbdRT-nis-ori                 TAGAAATCATATGGACAATATTCTACACATATTTAATGCATGTCTTATGACAAGATTATTAATCATACTTAAGCAATTAATTGTGATTGAATATTCTTTT 5150
Cs_LG2_264961294-264969213    TAGAAATCATATGGACAATATTGTACACATATTTAATGCATGTTTCATGACAAGATTATTGATCATACTTAAGCAATTAATTATGATTGAATATTCTTTC 5132
Cn_utg2999_44585-52528        TAGAAATCATATGGACAATATTCTACACATATTTAATGCATGTCTTATGACAAGATTATTAATCATACTTAAGCAATTAATTGTGATTGAATATTCTTTT 5129
Cs_LG3_149169260-149177129    TAGAAATCATATGGACAATATTGTACACATATTTAATGCATGTTTTATGACAAGATTATTAATCATA---------TTAATTGTGATTGAACATTCTTTC 5048
Cn_utg57260_32948-40788       TAGAAATCATATGGACAATATTGTACACATATTTAATGCATGTTTTATGACAAGATTATTAATCATA---------TTAATTGTGATTGAACATTCTTTC 5021
                                     5,220               5,240               5,260               5,280               5,300
SbdRT-nis-ori                 CAAGACACATTAAATATATAAAGATGATAATTGTCATACTAAATTTCATTAATATGAAATATGACAATGGAATATCAAAACTCTTTCTTGA-GAAATTAC 5249
Cs_LG2_264961294-264969213    TTAGACACATTAAATATATAAAGATGA-------------------------------------CAATGGAATATCAAAACTCTTTCTTGA-GAAATTAC 5194
Cn_utg2999_44585-52528        CTAGACACATTAAATATATAAAGATGATAATTGTCATACTTGTTTTCATTAATATGAAATATGACAATGGAATATCAAAACTCTTTCTTGA-GAAATTAC 5228
Cs_LG3_149169260-149177129    CTAGACACATTAAATATATAAAGATGATAATTGTCATACTTGTTTTCATTAATATGAAATATGACAATGGAATATCAAAATTCTTTTTTGAAGAAATTAC 5148
Cn_utg57260_32948-40788       CTAAACACATTAAATATATAAAGATGATAATTGTCATACTTGTTTTCATTAATATGAAATATGACAATGGGATATCAAAATTCTTTTTTGAAGAAATTAC 5121
                                     5,320               5,340               5,360               5,380               5,400
SbdRT-nis-ori                 TTGAGCGAGGACGGTAAGTAATTGAAAGGTTTTTTGAATTCATATAATCTTTGGAAGTGTACAAAGTGCAATAATTCATGAATGGATAAATGCATGCATC 5349
Cs_LG2_264961294-264969213    TTGAGCGAGGACGGTAAGTAATCGAAAGATTTTT-GAATTCATATAATCTTTGGAAGTGTACGAAGTGCAATAATTAATGAATGGATAAATGCATGCATC 5293
Cn_utg2999_44585-52528        TTGAGCGAGGACGGTAAGTAATCGAAAGGTTTTTTGAATTCATATAATCTTTGGAAGTGTACGAAGTGCAATAATTCATGAATGGATAAATGCATGCATC 5328
Cs_LG3_149169260-149177129    TTGAGCGAGGACGATAAGTAATTGAAAGGTTTTTTGAATTCATATAATCTTTGAAAGTGTACGAAGTGCAATAATTCATGAATGGATAAATGCATGCATC 5248
Cn_utg57260_32948-40788       TTGAGCGAGGACGATAAGTAATTGAAAGGTTTTTTGAATTCATATAATCTTTGAAAGTGTACGAAGTGCAATAATTAATGAATGGATAAATGCATGCATC 5221
                                     5,420               5,440               5,460               5,480               5,500
SbdRT-nis-ori                 TATGAAATTATTCTTATATTAAATTATTAAATCC-ATTTTCGATTTATAATTAAAATAAGATAAATTTTAAGTATTCCTGAAAGCTAGTGAGAGTATTAT 5448
Cs_LG2_264961294-264969213    TATAAAATTATTCTTATATTTAATTATTAAATCTTAGTTTTGATTTATAATTGAAATAAGATAAATTTTAAGTATTCCTAAAAGCTAGTGGGAGTATTAT 5393
Cn_utg2999_44585-52528        TATGAAATTATTCTTATGTTAAATTATTAAATCC-ATTTTTGATTTATAATTAAAATAAGATAAATTTTAAGTATTCCTGAAAGCTAGTGGGAGTATTAT 5427
Cs_LG3_149169260-149177129    TATGAAATTATTCTTATGTTTAATTATTAAATCCCATTTTCGATTTATAATTAAAATAAGATAAATTTTAAGTATTCCTGAAAGCTAGTGGGAGTATTAT 5348
Cn_utg57260_32948-40788       TATGAAATTATTCTTATGTTTAATTATTAAATCCCATTTTCGATTTATAATTAAAATAAGATAAATTTTAAGTATTCCTGAAAGCTAGTGGGAGTATTAT 5321
                                     5,520               5,540               5,560               5,580               5,600
SbdRT-nis-ori                 GTTAAGATT-TTAAATCTTATATGTTGAATAACATATAGGAAGTAACATTCATTGTAAGCTAAATGTTAAGTCTTAGATTGGAATGAAACTTCACTTATA 5547
Cs_LG2_264961294-264969213    GTTAAGGTT-TTAAATCTTATATGTTGTATAACATATGATAAGTAACATTCCTTGAAAGCTAAATGTTAAGTCTTAGATTGGAATGAAACTTCACTTGTA 5492
Cn_utg2999_44585-52528        GTTAAGGTT-TTAAATCTTATATTTTGTATAACATATGAGAAGTAACATTCCTTGAAAGCTAAATGTTAAGTCTTAGATTGGAATGAAACTTCACTTGTA 5526
Cs_LG3_149169260-149177129    GTTAAGTTTCTTAAATCTTATATGTTGTATAACATATAGTAAGTAACATTCCTTGTAAGCTAAATGTTATGTCTTAGATTGTAATGAAACTTCACTTGTA 5448
Cn_utg57260_32948-40788       GTTAAGTTTCTTAAATCTTATATGTTGTATAACATATAGTAAGTAACATTCCTTGTAAGCTAAATGTTATGTCTTAGATTGTAATGAAACTTCACTTGTA 5421
                                     5,620               5,640               5,660               5,680               5,700
SbdRT-nis-ori                 AGATTTAAGAATTAAAACATATAACCTTGAAATATATGTTTATTGTTAATAAACATGTGCAGTAAAA-TAAATGATTCTAAATAAAACATGATATATCAT 5646
Cs_LG2_264961294-264969213    AGATTTAAGAATTAAAACATACAACCTTGGAATATATGTTTATTATTAATAAACATGTGCAGTAAAAATAAATGATTCTAA-TAAAACATGATATATCAT 5591
Cn_utg2999_44585-52528        AGATTTAAGAATTAAAACATACAACCTTGGAATATATGTTTATTATTAATAAACATGTGCAGTAAAATTAAATGATTCTAA-TAAAACATGATATATCAT 5625
Cs_LG3_149169260-149177129    AGATTTAAGAATTAAAACATACAACCTTGGAATATATGTTTATTGTTAATAAACATGTGCAGTAAAA-TAAATGATTCTAA-TAAAACATGATATATCAT 5546
Cn_utg57260_32948-40788       AGATTTAAGAATTAAAACATACAACCTTGGAATATATGTTTATTGTTAATAAACATGTGCAGAAAAA-TAAATGATTCTAA-TAAAACATGATATATCAT 5519
                                     5,720               5,740               5,760               5,780               5,800
SbdRT-nis-ori                 TTGGGAAATTGATAAATATCTTTTGACAAGATAGAACGTGAATGAATTCTAAGGTTGACCTGTAGATTCATCATGTACATCATGTAGTGTATTAAAATTG 5746
Cs_LG2_264961294-264969213    TTGGGAAATGGATAAATATTTTTTGACAAGATAGAACGTGAATGAAGTCTAAGTTTGACCTGTAGATTCATCATGTACATCATGTAGTGTATTTAAATTG 5691
Cn_utg2999_44585-52528        TTGGGAAATGGATAAATATTTTTTGACAAGATAGAACGTGAATGAAGTCTAAGTTTGACTTGTAGATTCATCATGTACATCATGTTGTGTATTAAAATTG 5725
Cs_LG3_149169260-149177129    TTGGGAAATTGATAAATTTCTTTTGACAAGATAGAACGTGAATGAATTCTAAGGTTGACCTGTAGATTCATCATGTACATCATGTTGTGTATTAGAATTG 5646
Cn_utg57260_32948-40788       TTGGGAAATTGATAAATATCTTTTGACAAGATAGAACGTGAATGAATTCTAAGGTTGACCTGTAGATTCATCATGTACATCATGTTGTGTATTAGAATTG 5619
                                     5,820               5,840               5,860               5,880               5,900
SbdRT-nis-ori                 TTGTATCATTCTTAAATGATAGATGATTTGACAACATTGATATCTAATTAAAAGTTATTAGATAACTTGTGTAATGAATATTCTAATAGAATTACATCCT 5846
Cs_LG2_264961294-264969213    TTGTATCATTCTTAAATAAGAAGTGATTTGACAACATTGATATCTAATTAAAAGTTATTAGATAACTTGTGTAATGAATAGTCTAATAGAATTACATCCT 5791
Cn_utg2999_44585-52528        TTGTATCATTCTTGAATAAGAAGTGATTTGACAACATTGATATCTAACTAAAAGTTATTAGATAACTTGTGTAATGAATATTCTAATAGAATTACATCCT 5825
Cs_LG3_149169260-149177129    TTGTATCATTCTTGAATAAGAAGTGATTTGACAACATTGATATCTAACTAAAAGTTATTAGATAACTTGTGTAATGAATATTCTGAAAGAATTACATCCT 5746
Cn_utg57260_32948-40788       TTGTATCATTCTTGAATAAGAAGTGATTTGACAACATTGATATCTAACTAAAAGTTATTAGATAACTTGTGTAATGAATATTCTCAAAGAATTACATCCT 5719
                                     5,920               5,940               5,960               5,980               6,000
SbdRT-nis-ori                 AAAA-CCTGGAATTGCTCCTAGATTCTATGGATGAACAATATGTCAAATGTTGTGCAAATTGTCATCAGCTTTAAACTTGAAAGTTAGGAAAGCATCAAA 5945
Cs_LG2_264961294-264969213    TAAAACCTGGAATTGCTCCTAGATTCTATGGATGAACAATATGTCAATTGTTGTGCAAATTGTCATCAGCTTCAAACTTGAAAGTTTGGAAGACATCAAA 5891
Cn_utg2999_44585-52528        CAG--CCTGGAATTGCTCCTAGATTCTATGGATGAACAATATGTCAAATGTTGTGCAAATTGTCATCAGCTTTAAACTTGAAAGTTAGGAAAGCATCAAA 5923
Cs_LG3_149169260-149177129    CAG--CCTGGAATTGCTCCTAGATTCTATGGATGAACAATATGTCAAATGTTGTGCAAATTGTCATCAGCTTTAAACTTGAAAGTTAGGAAAGCATCAAA 5844
Cn_utg57260_32948-40788       CAA--CCTGGAATTGCTCCTAGATTCTATGGATGAACAATATGTCAAATGTTGTGCAAATTGTCATCAACTTTAAACTTGAAAGTTAGGAAAGCATCAAA 5817
```

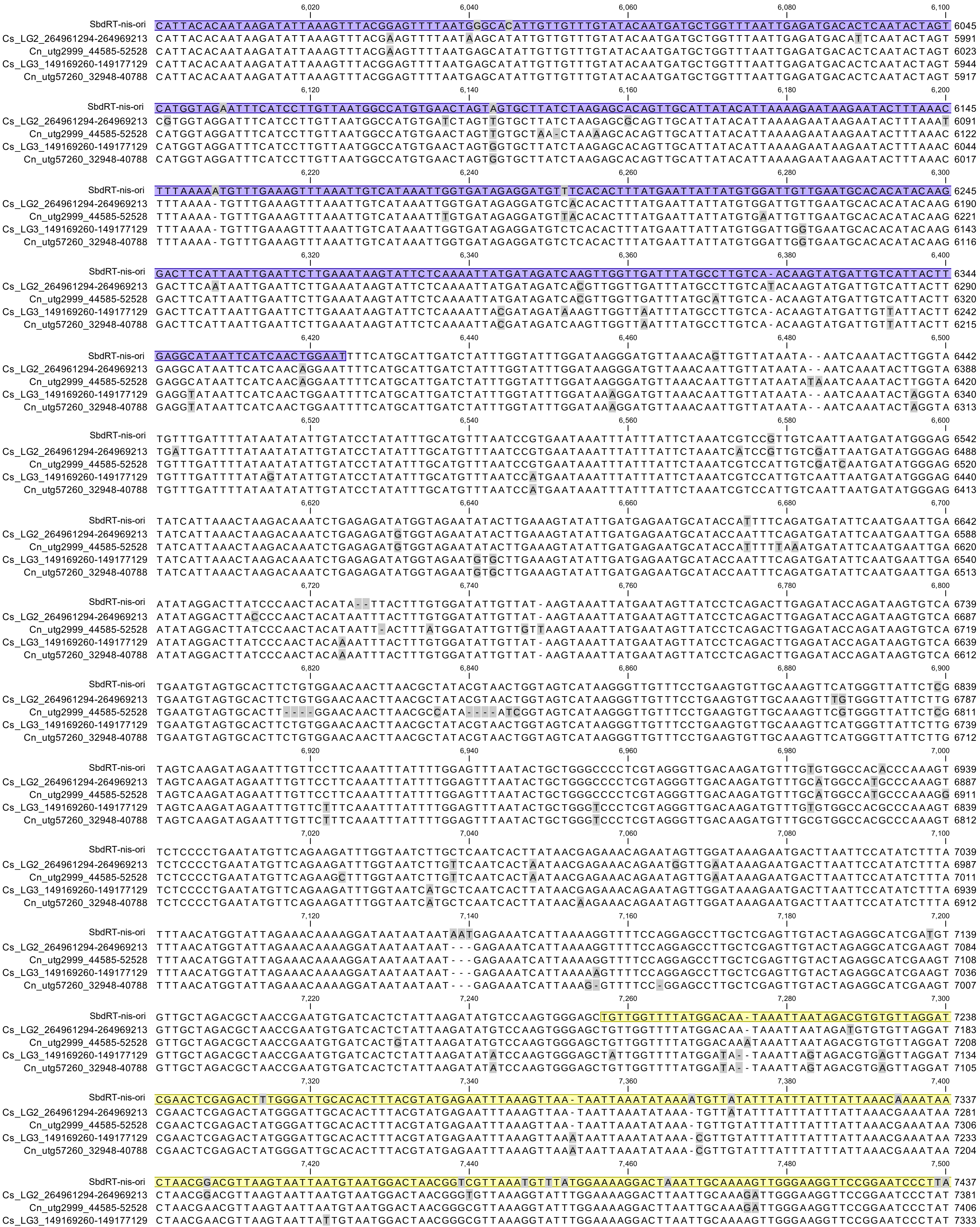
SbdRT-nis-ori
Cs_LG2_264961294-264969213
Cn_utg2999_44585-52528
Cs_LG3_149169260-149177129
Cn_utg57260_32948-40788

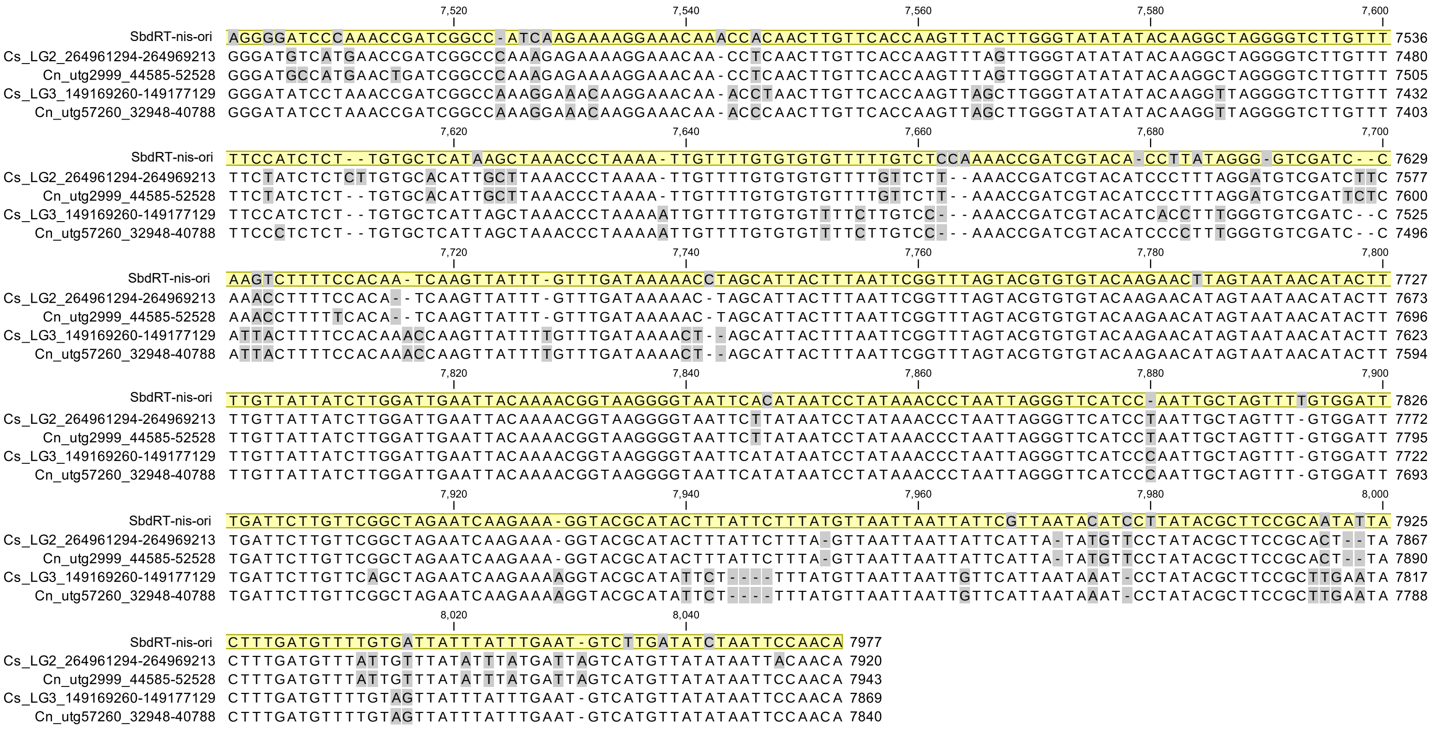

S**upplementary Figure. 5 | The structures of SbdRTs in *C. seticuspe* and *C. nankingense.* a**, DNA sequences of SbdRT-nis-type copies in *C. seticuspe* and *C. nankingense* are aligned. Yellow, pink, and blue boxes represent LTR, PBS-ATG, and NIS, respectively. Cs_LG2_264961294-264969213 and Cs_LG3_1449169269-1449177129 are SbdRT-nis copies in *C. seticuspe.* Cn_utg2999_44585-52528 and Cn_utg57260_32948-40788 are SbdRT-nis copies in *C. nankingense*. Gray regions show sequence differences.

b

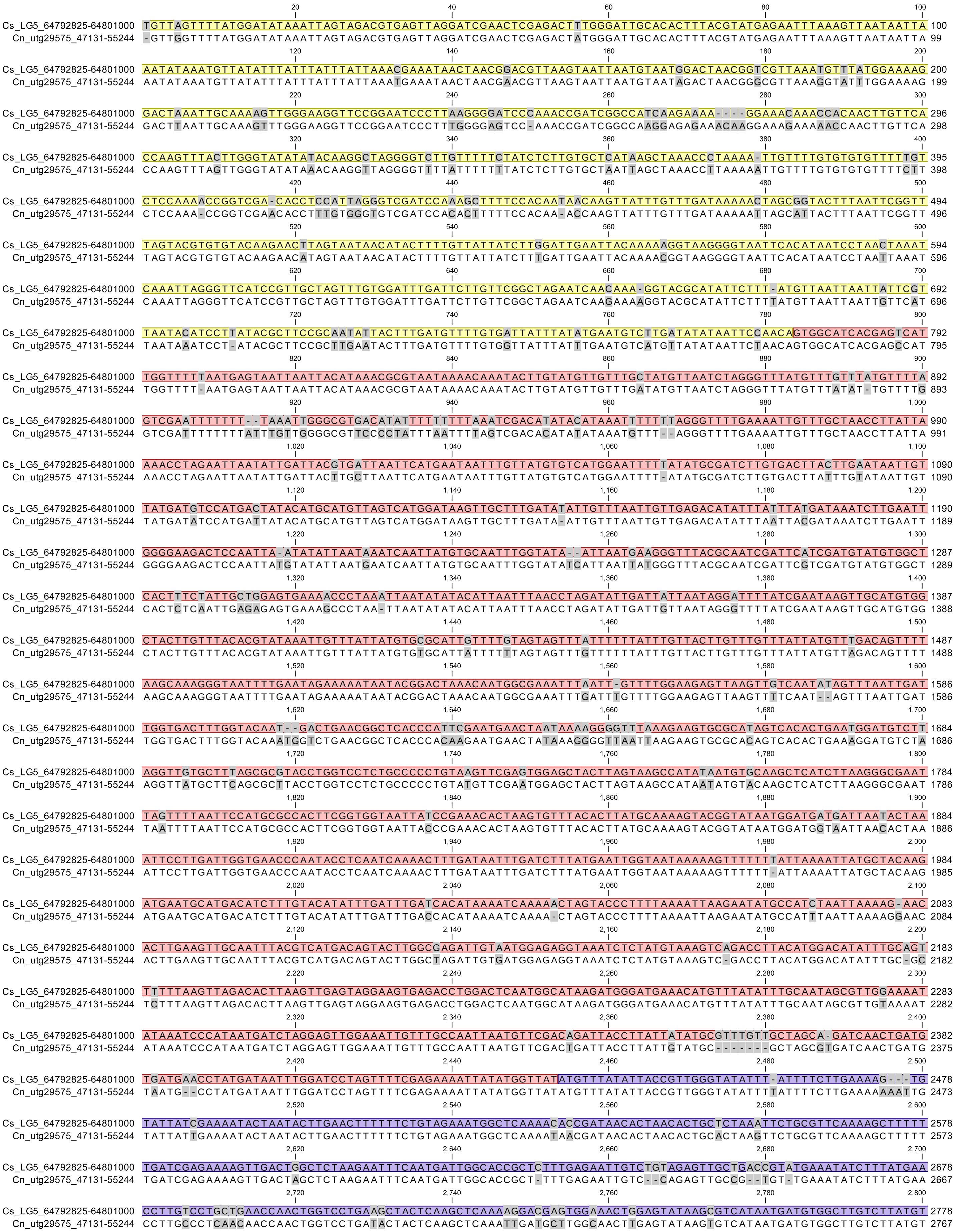

```
                           2,820 2,840 2,860 2,880 2,900
Cs_LG5_64792825-64801000   TGGGAACAATGGAATCTGCTCTTCAAAAGCAGTTCGAACATTATTTCCCATATAATATGCTCAATGAACTTCGAACTATGTTTGAGAAACCAGCTGCCGT 2878
Cn_utg29575_47131-55244    TGGGAACAATGGAATCTACTCTTCAAAAGCAGTTATAACATTATTTCCCATATAATATGCTCAATGAACTTCGGACTATGTTTGAGAAACCAGCTGCCGT 2867
                           2,920 2,940 2,960 2,980 3,000
Cs_LG5_64792825-64801000   TGAGATTTATGACTTGACCGACGCGTTACACAGCTGTAGGCAGGGAGAGGGTAAACCCGTAAGTGCTCATGTGCTGGAAATGAAAGGTTACATGGACCAG 2978
Cn_utg29575_47131-55244    CGAGATTTATGACTTGACCGACACGTTACACAGCTGTAGGCAAGGTGAGGGTAAACCCGTAAGTGCTCATGTACTGGAAATGAAAGGTTACATGGACCAG 2967
                           3,020 3,040 3,060 3,080 3,100
Cs_LG5_64792825-64801000   CTAAAAGCACTTGGCAAAACTTATGAGGATGACATGGCTGTGAATCTGATTAATCGATCCCTTAACAAGGATTTTAATGGTTTTGTCAGAAATTTCAATA 3078
Cn_utg29575_47131-55244    CTCAAAGCACTTGGCAAAACTTATGATGATGACATGGCTGTGAATCTGATTAATCGATCCCTTAACAAGGATTTTAATGATTTTGTCAGGAATTTCAATA 3067
                           3,120 3,140 3,160 3,180 3,200
Cs_LG5_64792825-64801000   TGCACTGCGTGGGGAAGACAGTTACTGACCTACACGCATTGCTAATTGACTATGAGAAAGGGTTGCCTCAAAAGGTTCCTACCGCCCAGGTACTTGCTAT 3178
Cn_utg29575_47131-55244    TACAC--CGCTGGGAAGACAAGCATCGACCTACACGCATTGCTTATTGATTATGAGAAAGGGTTGCCTCAAAAGGTTCCTACTGCCCAGGTACTTGCTAT 3165
                           3,220 3,240 3,260 3,280 3,300
Cs_LG5_64792825-64801000   TCAGGGTGGAAGAATTAATAAGCCAAAACCGCAGCGTGCGAACAACAAGAAGGGGAAGGGTAAGGGAAAGGTTGACAAAGGCAAACAGGTGATGGCCTAT 3278
Cn_utg29575_47131-55244    TCAGGGTGGAAGGATTAATAAGCCAAAACCGCATCGTGTGAACAACAAGAAGGGGAAGGGTAAGGGTAAGGTTGACCAAGGCAAACAGGTGATGGCCTAT 3265
                           3,320 3,340 3,360 3,380 3,400
Cs_LG5_64792825-64801000   CAACCTCAACCCAAGAAGAACCCTACTCCTAGGAAGGATAATCCTAAAAAGGATCAAGCATGCCACCATTGCAAGACAGTTGGGCATTGGAAGAGAAACT 3378
Cn_utg29575_47131-55244    CAACCTCAACCCAAGAAGAACCCTACTCCTAGGAAGTAAAATCCTAAAAAGGATCAAGCATGCCACCACGGTAAGATCGTTGGGAATTGGAAGAGAAACT 3365
                           3,420 3,440 3,460 3,480 3,500
Cs_LG5_64792825-64801000   GTGCTCTCTATCTTGATGAATTGCGGACAAATAAGATGAA---TGCTGGGTCCTCGGGCGCCGGTTCTTCAGGTAATTTATTTGTCATTGAATTATTTAC 3475
Cn_utg29575_47131-55244    GTGCTCTCTATCTTGATGAATTGCGGGCAAATAAGAAGAAGTATGAAGGGTCCTTGGGCGCCGGTTCTTCTGGTAATTTATTTGTCATTGAATTATTTAC 3465
                           3,520 3,540 3,560 3,580 3,600
Cs_LG5_64792825-64801000   TCTAAATCATAATAATTCCTGGGTATATGATACGGGCTG----------TGGTATTCACATTTGTAATACACTGCAGGGGTTTAGAGTGGAAAGGAAGCT 3565
Cn_utg29575_47131-55244    TCTAAATCATAATAATTCCTGGGTATATGATACGGGCTGATACGGGCTGTGGTATTCACATTTGTAATACACTGCAGGGGTTTAGAGTGGAAATGAAGCT 3565
                           3,620 3,640 3,660 3,680 3,700
Cs_LG5_64792825-64801000   TGCCTATGGGGAGCAATATCTGCACGTGGGAAATGGCGCACATGCAGCTGTTGAAGCCATAGGAACGTTTAGTTTAGTATTGCCTTCAGGCTTAGTACTG 3665
Cn_utg29575_47131-55244    TGCCTATGGGGAGCAATATCTG-ACGCGGGAAATGGCGCATGCACAA-----GAAGCCATAGGAACGCTTAGTTTAGTATTGCCTTCGGGCTTAGTATCG 3659
                           3,720 3,740 3,760 3,780 3,800
Cs_LG5_64792825-64801000   TTATTGAACAATTGTCATTACGCACCTTCTATAATTAGAGGTGTTGTTTCCTTTTCAAGTTTGTTGGATGATGGTTTTACTCATACTATTTCAAGTAATG 3765
Cn_utg29575_47131-55244    TTATTAAACAATTGTCATTACACACCTTCTATAATTAGAGGTGTTGTTTCCTTTC--GCTTTGTTGGATGATGGTTTTACTCATACTATTTCACGTAATG 3757
                           3,820 3,840 3,860 3,880 3,900
Cs_LG5_64792825-64801000   GAATTGATGTTTCTTTAAATGGTGTATTTTATTTTAAAGCAATTACTGTGAATGGTGTGTTTGAAATTGATATGATTGGAAATGTATCTAACAATAATAA 3865
Cn_utg29575_47131-55244    GAATTGATGTTTCTTTAAATGGTGTATTTTATTTTAAAGCAATTACTGTGAATGGTGTGTTTGAAATTGATGTGATTGGAAATGAATCTAACAATAATAA 3857
                           3,920 3,940 3,960 3,980 4,000
Cs_LG5_64792825-64801000   TAATTCAATGTTTAGCATTAACAAGAAGATAAAGTTAGACTTGAATTCTTCTTATCTTTGGCATTGTCGACTTGCTCATATAGGCAAGGCTCGCATGCAT 3965
Cn_utg29575_47131-55244    TAATTCGATGTTTAGCATTAACAAGAAGATACAGTTAGACTTGAATTCTTCTTATCTTTGGCATTGTAGACTTGCTCATATTGGCAAGGCTCGCATGCAT 3957
                           4,020 4,040 4,060 4,080 4,100
Cs_LG5_64792825-64801000   AAGCTACAACAAGAAGGGCTTTTGCAAAATATCAATGATGAATCATTTGATAAATGTGAGTCATGCATATCTGGGAAGATGACTAAGAAACCTTTTCATA 4065
Cn_utg29575_47131-55244    AAGCTGCAACAAGAAGGGCTTTTGCAAAATATCAATGATGAATCATTTGATAAATGTGAGTCATGCATATCTGGGAAGATGACGAAGAAACCTTTTCATA 4057
                           4,120 4,140 4,160 4,180 4,200
Cs_LG5_64792825-64801000   GCAATATGGAGAGGGCTTCTGATCTACTTGGTCTTATTCATACTGATGTGTGTGGACCACTTAGACATGTGTCAAAGAGAGGTGCTAGCTACTTCTTAAC 4165
Cn_utg29575_47131-55244    GCAATATGGAGAGGGCTTCTGATCTACTTGGTCTTATTCATACGGATGTGTGTGGACCACTTAGACATGTGTCAAAGAGAGGTGCTAGCTACTTCTTAAC 4157
                           4,220 4,240 4,260 4,280 4,300
Cs_LG5_64792825-64801000   CTTTACTGATGATTTCAGCCGTTATGGTTATGTTTATTTACTTAAGCATAAACATGAGGTGTTTGAAACATTTAAGGTTTTCAAAGCTGAAGTAGAACTG 4265
Cn_utg29575_47131-55244    CTTTACTGATGATTTCAGCCGTTATGGTTATGTTTATTTACTTAAGCATAAACATGAGGTGTTTGAAACATTTAAGGTTTTCAAAGCTGAAGTAGAACTG 4257
                           4,320 4,340 4,360 4,380 4,400
Cs_LG5_64792825-64801000   CAATTGGGAAAGAAGATTAAAGCTCTTCGATCCGATAGAGGAGGCGAGTATCTAAGCCAAGAGTTTAAAGATTACTTAGCTGAGAATGGAATTGTCCAGA 4365
Cn_utg29575_47131-55244    CAATTGGGAAAGGAGATTAAAGCTCTTCGATCCGATAGAGGA----AGTATCTAAGCTAAGAGTTTAAAGATTACTTAGTCGAGAATGGAATTGTCCAGA 4353
                           4,420 4,440 4,460 4,480 4,500
Cs_LG5_64792825-64801000   ATCTCACTTCTCCATATACTCCACAACAAAATGGTGTATCTGAGAGAAGAAATCGTACATTGTTGGACATGATACGATCTATGTTTAATCTTACTACTCT 4465
Cn_utg29575_47131-55244    ATCTCACTTCTCCATATACTCCACAACAAAATGGTGTATCTGAGAGAAGAAATCGTACATTGTTGGACATGATACGATCTATGTTTAATCTTACCACTCT 4453
                           4,520 4,540 4,560 4,580 4,600
Cs_LG5_64792825-64801000   ACCTTTATCCTTTTGGGATTATGCTTTAGAATCTGCTGTTCGTATTCTAAATATGGTTCCCACAAAGAAGGTTGATAAGACACCATATGAAATATGGCAT 4565
Cn_utg29575_47131-55244    TCCTTTATCCTTTTGGGATTATGCTTTGGAATCAGCTGTTCGCATTCTAAATATGGTTCCCACAAAGAAGGTTGATAAGACACCATATGAAATATGGCAT 4553
                           4,620 4,640 4,660 4,680 4,700
Cs_LG5_64792825-64801000   GGTAAAGCTCCTAACCTGTCTTATTTAAAGGTCTGGGGTTGCGAGGCGTATGTTAGACGTGACTCAGCAGATAAACTCCAACAAAGATCTGTAAAGTGTA 4665
Cn_utg29575_47131-55244    GGTAAAGCTCCTAACCTGTCTTATTTAAAGGTCTGGGGTTGCGAGGCGTATGTTAGACGTGACTCAGCAGATAAACTCCAACAAAGATCTCAGAAGTGTA 4653
                           4,720 4,740 4,760 4,780 4,800
Cs_LG5_64792825-64801000   TCTTCGTAGGATACCCTAAGGAAACTATGGGTTACTACTTTTACTATCCAACTGATAATAAAGTAATTGTTGCTAGGTATGGAGATTTTCTTGAAAGAGA 4765
Cn_utg29575_47131-55244    TCTTC--AGGATACCCTAAGGAAACTTTGGGTTACTACTTTAAG-ATCCAACCGAAAATAAAGTAATTGTTGCTAGGTATGGAGATTTTCTTGAAAGAGA 4750
                           4,820 4,840 4,860 4,880 4,900
Cs_LG5_64792825-64801000   TCTCGTATCTCAAGAATTCAGTGGGAGAGTTTATGATCTGGAAGATGATCATGAAGATGCATTACCTTCTGAAAACACTAACGTACT-TCCGGTAGAACC 4864
Cn_utg29575_47131-55244    TCTCGTATCTCAAGAAT----TGGGAGAGTTTATGATCCGGAAGATGATCAAGAAGATACATTACCTTC--AAAACACTAACGCATTCTCCGGTAGAACC 4844
                           4,920 4,940 4,960 4,980 5,000
Cs_LG5_64792825-64801000   TGATAGTCTGGGTCCTCCACCAGAAGTTGAAGTGATTCCCGTTCGTAGATCCGTAAGGAATATACGTGCACCTGAACGTCTATGTCTTAACGTAGAGGTT 4964
Cn_utg29575_47131-55244    -GATAGTCCGGGTCCTCCACCGTAAGTTGAAGTGATTCCG--TCAGAGATCCGTAAGGAATACGCGTGCACCTGAACGTCTATGTCTTAACGTAGAGGTT 4941
                           5,020 5,040 5,060 5,080 5,100
Cs_LG5_64792825-64801000   GAGGAAACTATGTTAGGAGATCTTGGAGAGCCTACTAGTTATAAAGCTGCCATGGCAAGTCCTGACAAGGCTAAATGGCAGGTTGCAATGAATGGGGAAA 5064
Cn_utg29575_47131-55244    GAGGAAACTATGTTAGGAGTTCTTGGAGAGCCTACTAGTTATAAAGCTGCCATGGCAAGTCCTGATAAGGATAAATGGCAAGTTGCAATGAATGGGGAAA 5041
                           5,120 5,140 5,160 5,180 5,200
Cs_LG5_64792825-64801000   TGCAATCCATGAGAGAAAATGAAGTCTGGGTAGAGGTTGACTTACCTCCAAACGCCTCAGTGGTTGGGAGTAAGTGGCTCTATAAGATTAAGACTGACAT 5164
Cn_utg29575_47131-55244    TGCAATCCATGAGAGAAAATGAAGTCTGGGTAGAGGTTGACTTACCTCCAAACGCCACGGTGGTTGGAAGTAAGTGGCTCTATAAGATTAAGACTGACAT 5141
                           5,220 5,240 5,260 5,280 5,300
Cs_LG5_64792825-64801000   GGATGGCAATGTACACGTCTATAAAGCTCGTCTAGTTGCGCAAGGTTTTACTCAAACTTATGGCATTGATTATGAGGAGACCTTTTCTCCT-GTAGCAGA 5263
Cn_utg29575_47131-55244    GGATGGCAATGTACATGTCTATAAAGCTCGTCTAGTTGCGCAAGGTTTTACTCAAACTTATGGCATTGATTACGAGGAGACCTTTTCTCCTCAGAATGGA 5241
                           5,320 5,340 5,360 5,380 5,400
Cs_LG5_64792825-64801000   CATTAGAGCTATTAGAATTCTCATAGCTATAGCAGCATACTATGACTATGAGATATGGCAAATGGATGTCAAAACCGCCTTCTTAAATGGTCGCTTAGAT 5363
Cn_utg29575_47131-55244    CATTAGAGCTATTAGAATTCTCATAGCTATAGCAG-ATACTATGACTATGAGATATGGCAAATAGATGTCAAAACCGC-TTCTTAAATGGTCGCT-AGAT 5338
                           5,420 5,440 5,460 5,480 5,500
Cs_LG5_64792825-64801000   GAAGACATCTATATGAAGCAACCTGAGGGGTTTGTGAATCCACAATTTCCAAACCGTGTATGCAAATTTCAAAGGTCCATTTATGGATTGAAGCAAGCAT 5463
Cn_utg29575_47131-55244    GAAGACATCTATATGAAGCAACCTGAGGGGTTCGTGAATCCACAATT---AAACCGTGTATGCAAACTTCAAAGGTCCATTTATGGATTGAAGCAAGCAT 5435
                           5,520 5,540 5,560 5,580 5,600
Cs_LG5_64792825-64801000   CAAGGCAATGGAATAAAAGATTTGATGAAGAAATCAAAAGGTATGGATTTGTTCAAAATCTTGATGAGCCATGTGTATATCACAAAGTTAGTGGGAGTGA 5563
Cn_utg29575_47131-55244    CAAGGCAATGGAATAAAAGATTTGATGAAGAAATCAAAAAGTATGGATTTATTCAAAATCTTGATGAGCCATGTGTATATCACAAAGTTAGTGGGAGTGA 5535
```

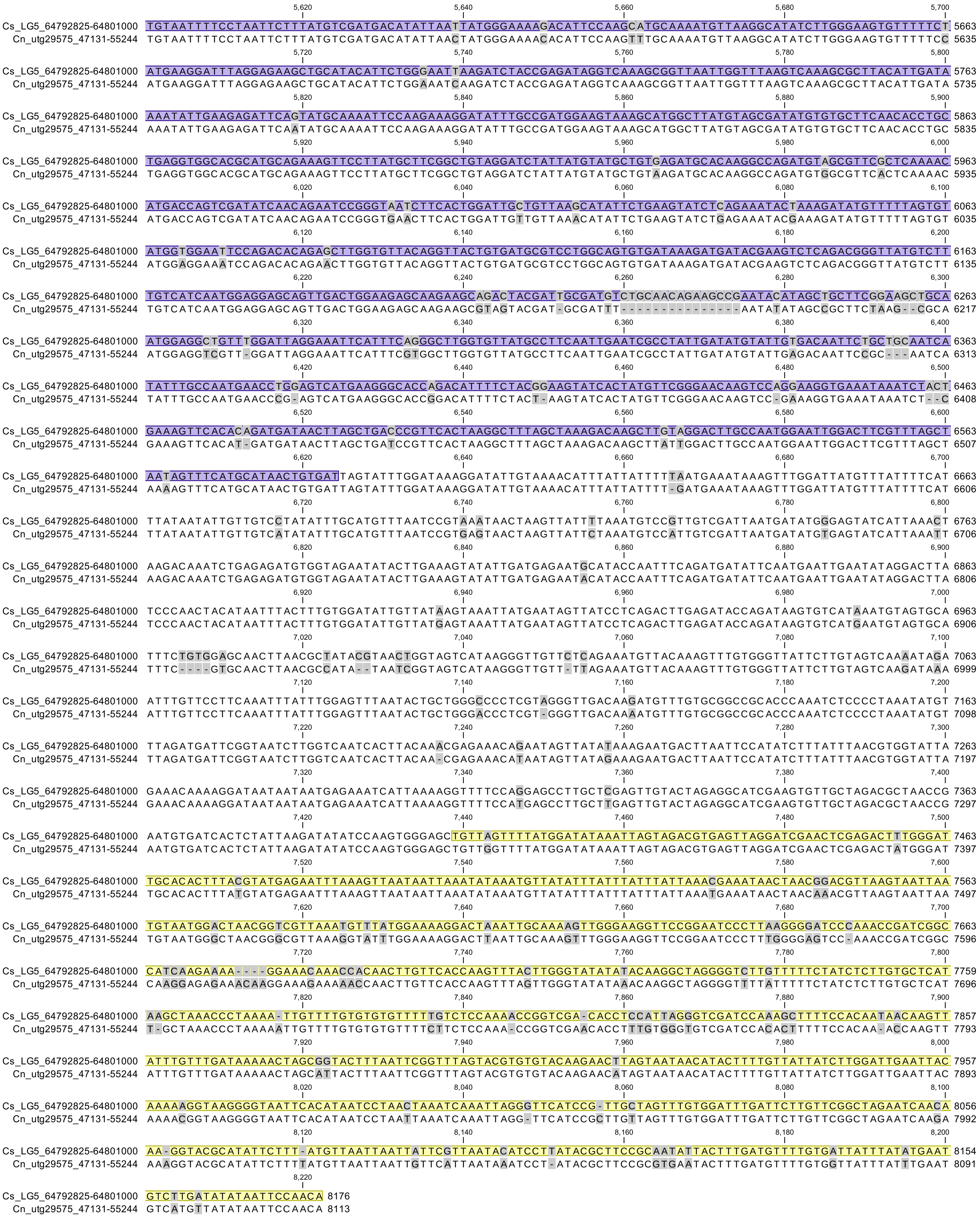


**b,** DNA sequences of SbdRT-orf-type copies in *C. seticuspe* and *C. nankingense* are aligned. Yellow, pink, and blue boxes indicate LTR, PBS-ATG, and ORF, respectively. Cs_LG5_64792825-64801000 and Cn_utg29575_47131-55244 are SbdRT-orf in *C. seticuspe* and *C. nankingense,* respectively.

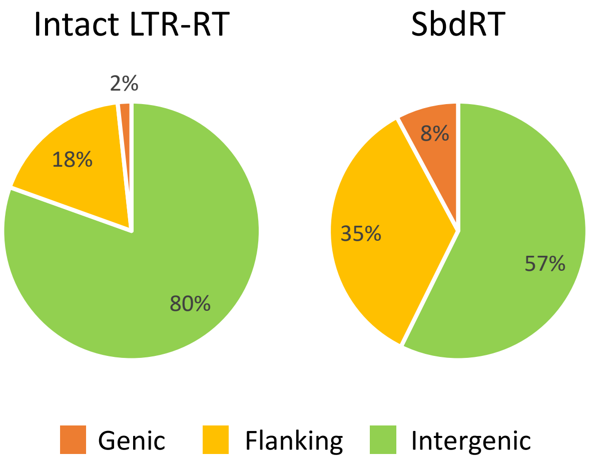


**Supplementary Figure 6 | Distribution of insertion sites of SbdRT in the *C. seticuspe* genome.** Genic, insertion of structural genes; flanking, insertions within 5 kb upstream or downstream of structural genes; intergenic, insertions in other regions. Intact LTR-RT represents all types of LTR-RT with LTRs at both ends. SbdRT represents all 360 SbdRT copies.

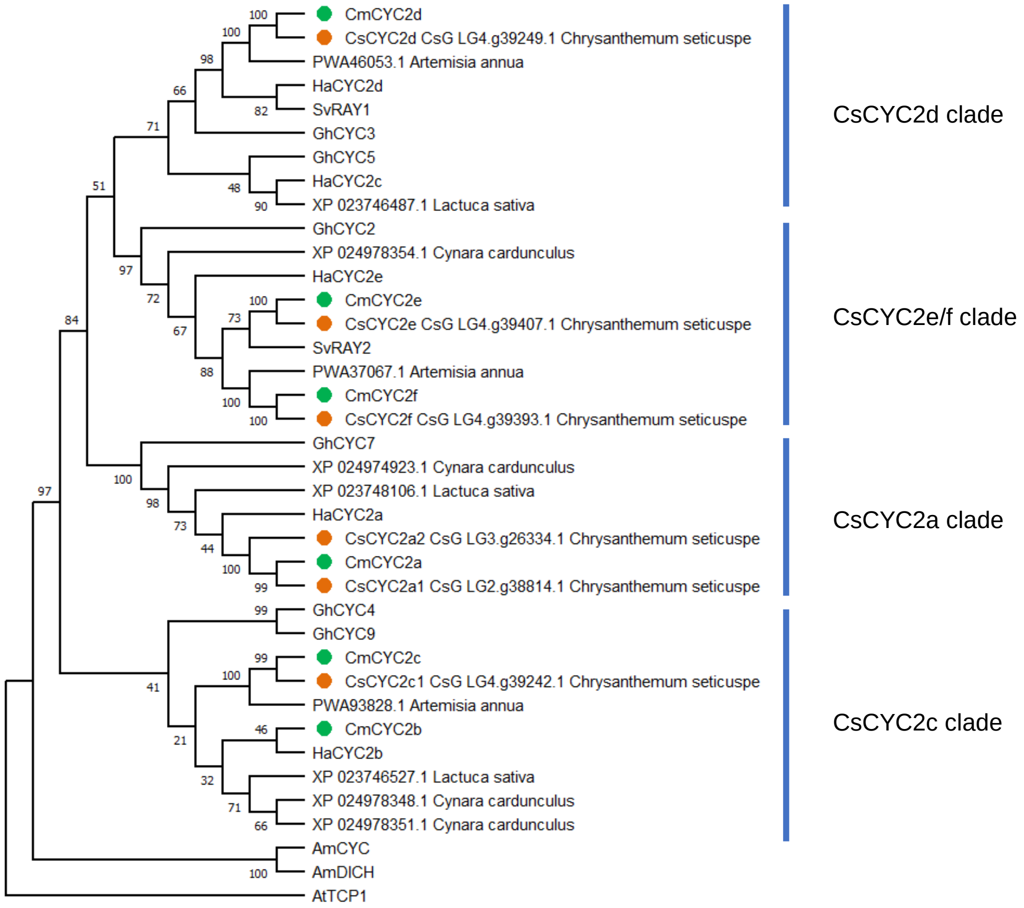


**Supplementary Figure 7 | Phylogenetic tree of the *CYC2* family genes in Asteraceae.** CsCYC2, CmCYC2, HaCYC2, GhCYC, and SvRAY are the *CYC2* family genes of *C. seticuspe*, *C. morifolium*, sunflower, gerbera, and *Senecio vulgaris,* respectively. The *CYC2* family genes of *C. seticuspe* and *C. morifolium* are marked with orange and green circles, respectively.

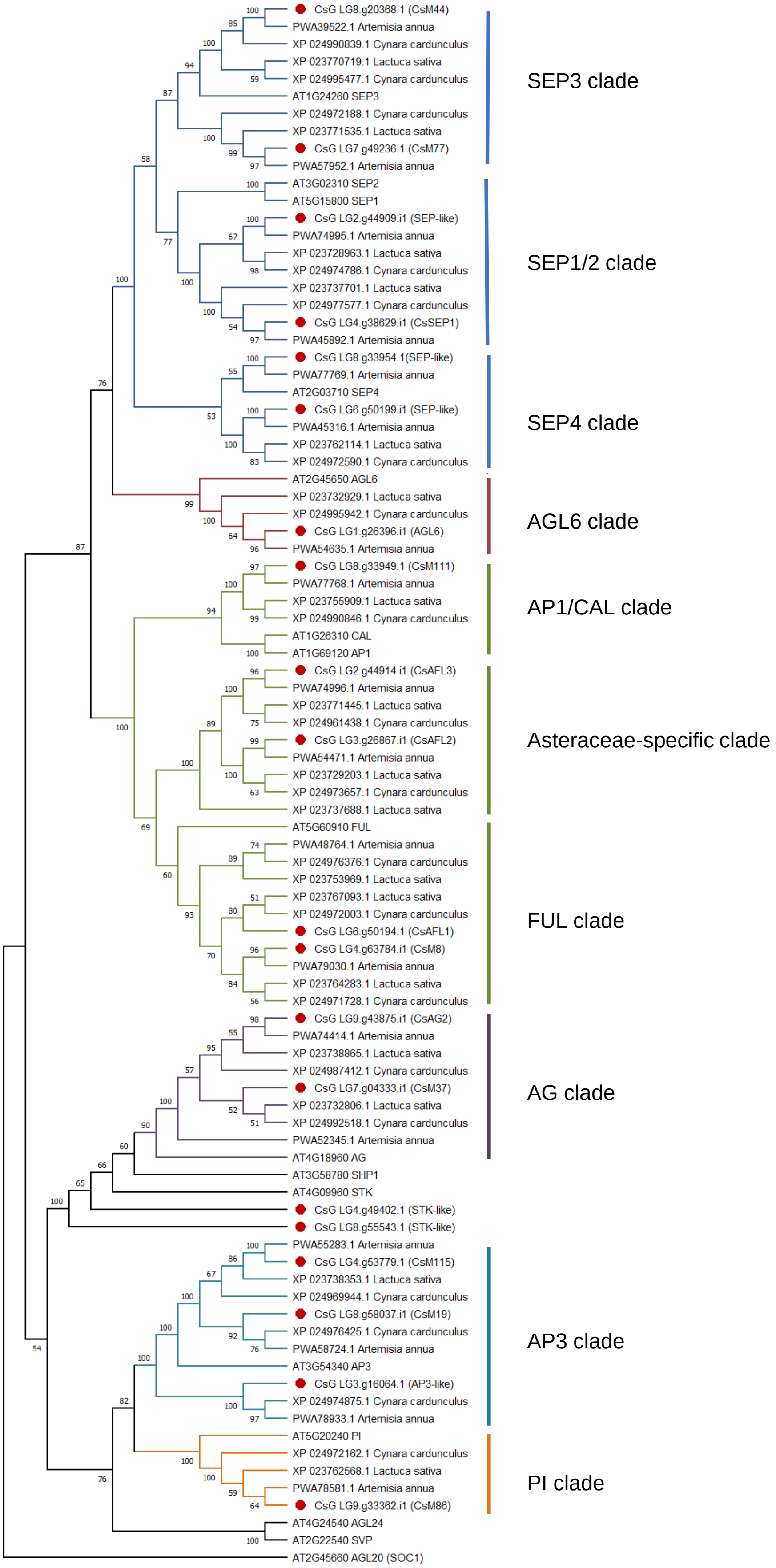


**Supplementary Figure 8 | Phylogenetic tree of ABCE MADS-box genes in Asteraceae.** MADS-box genes of *C. seticuspe* are marked with red circles.

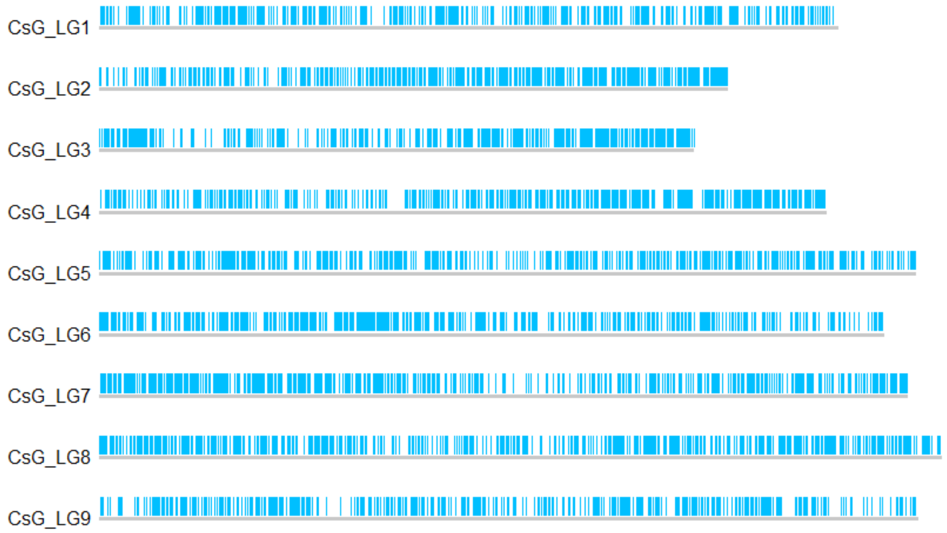


**Supplementary Figure 9 | Chromosomal locations of SSR markers in the *C. seticuspe* genome.** SSR markers detected in AEV02 were mapped to *C. seticuspe* pseudochromosomes.

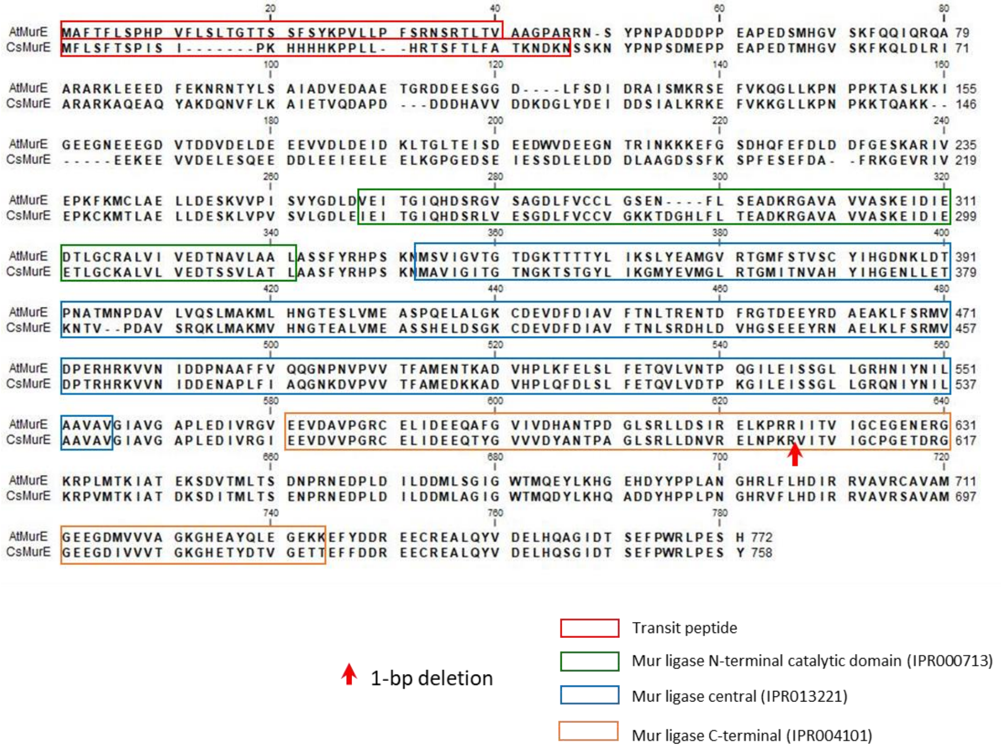


**Supplementary Figure 10 | Alignment of AtMurE and CsMurE proteins.** CsMurE retains chloroplast transit peptide and Mur ligase domains. The red arrow indicates the position of a single base pair deletion in *alb1*.

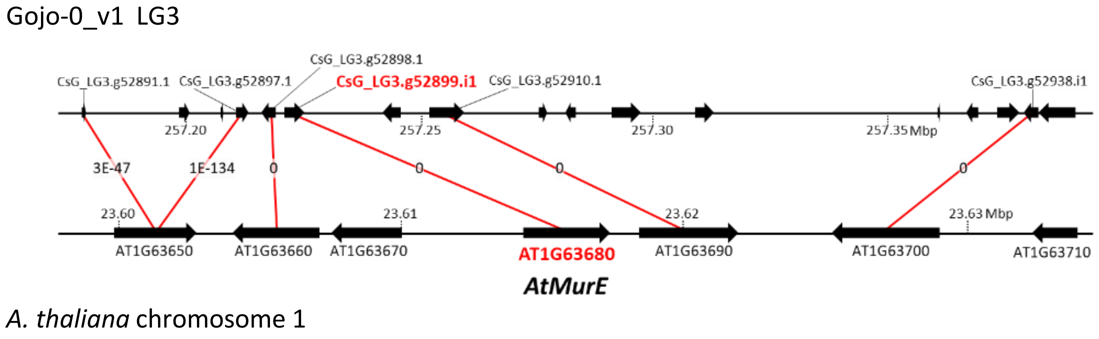


**Supplementary Figure 11 | Microsynteny between the *CsMurE* and *AtMurE* regions.** The predicted genes around LG3.g52899.i1 in *C. seticuspe* and *AtMurE* in *A. thaliana* are shown. The values shown on the red lines indicate e-values obtained from BLAST analysis.

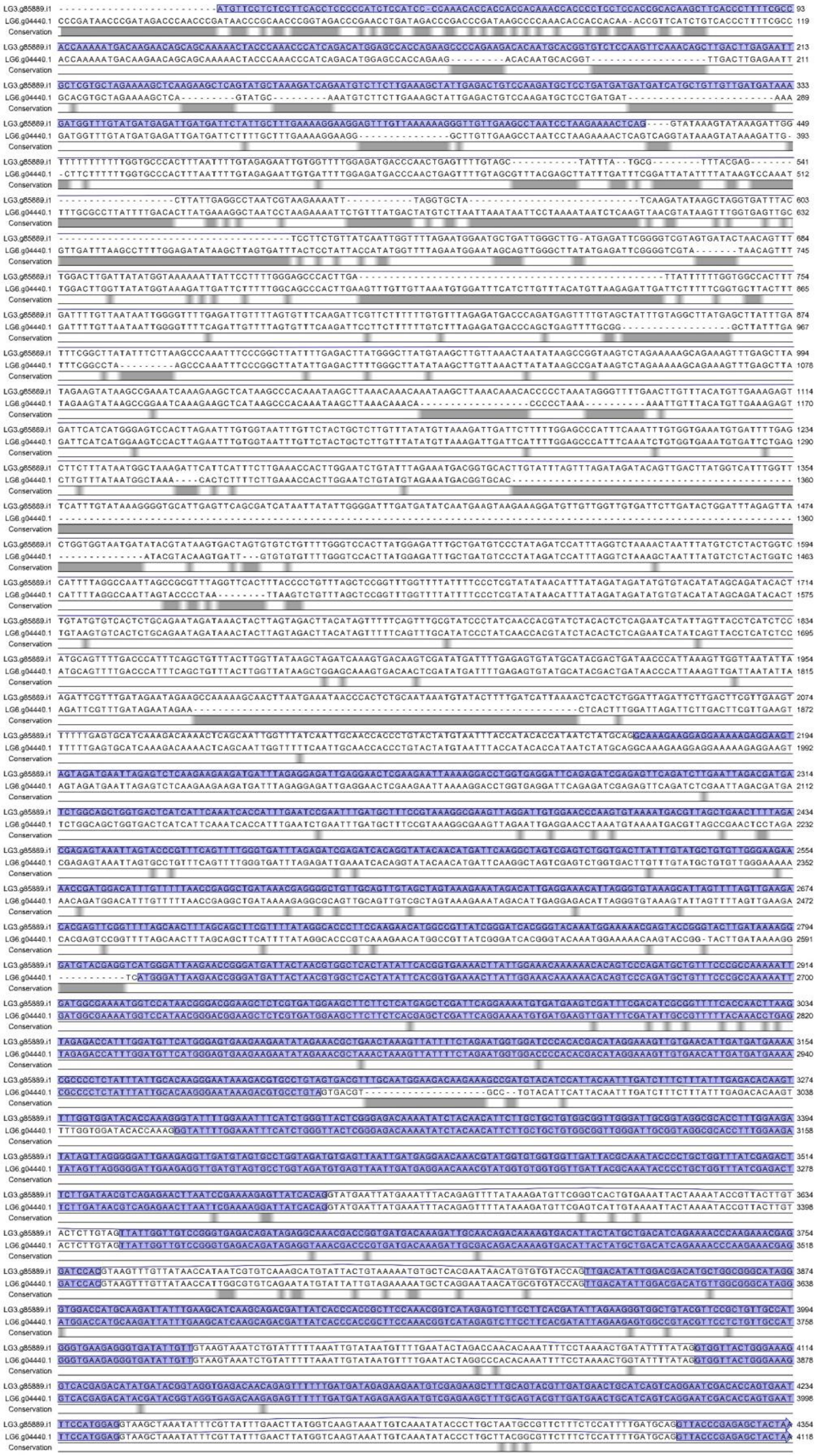

**Supplementary Figure 12 | Alignment of genome sequence of LG3.g85889.i1 (*CsMurE*) and LG6.g04440.1 (pseudo gene).** Purple regions are coding sequences. Gray regions show sequence differences between LG3.g85889.i1 and LG6.g04440.1. LG6.g04440.1 contains numerous deletions particularly in 5′ regions and does not encode a functional MurE.

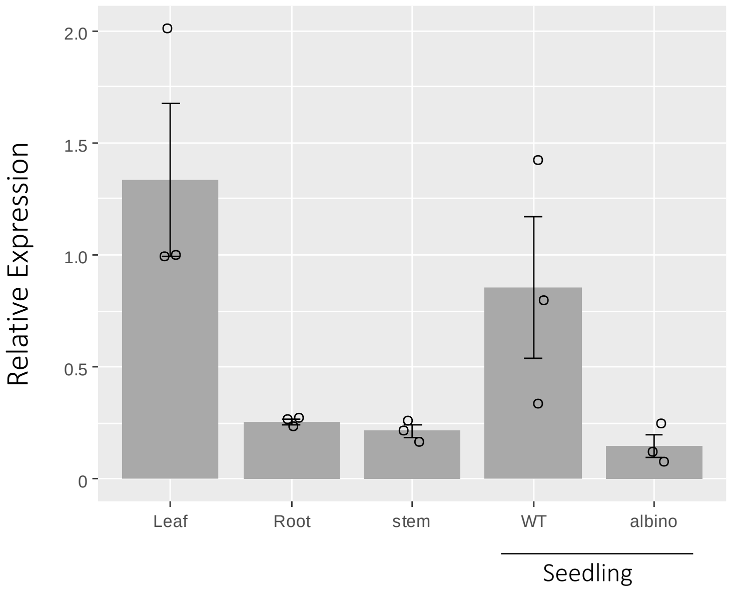


**Supplementary Figure 13 | Expression of *CsMurE* in various tissues.** Expression of *CsMurE* in wild-type (WT) and *alb1* was determined by qRT-PCR. *CsActin* was used as a reference. Data are presented as means ± standard errors (*n* = 3).

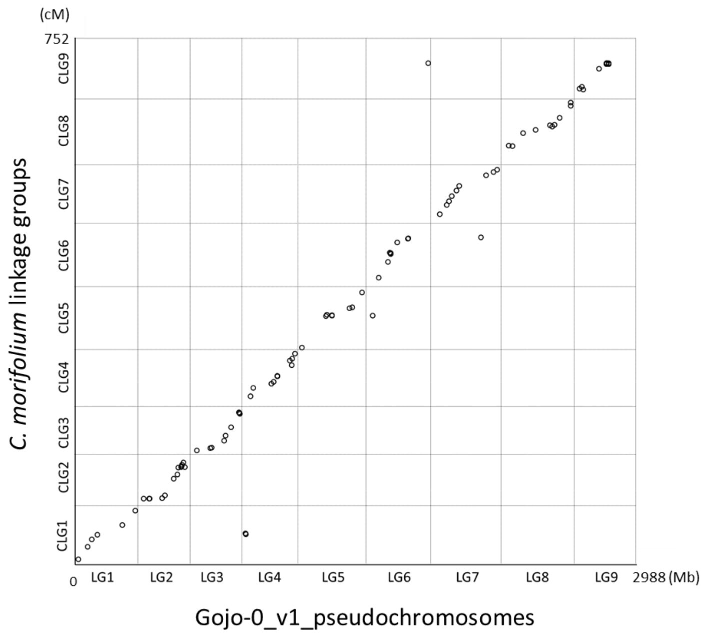


**Supplementary Figure 14 | Comparison of Gojo-0 genomic sequences with genetic map of *C. morifolium.*** The final assembly (Gojo-0_v1) was subjected to BLASTN search against 92 SNP markers of *C. morifolium*[1]. Markers showing significant hits (≤ 1e-10) were plotted on the pseudochromosomes of Gojo-0, and then linkage group names were assigned for individual chromosomes.

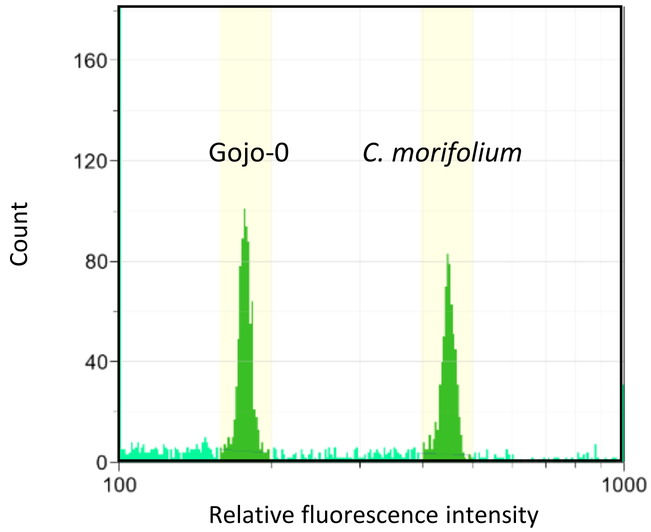


**Supplementary Figure 15 | Estimation of genome size of *C. morifolium* by ploidy analyzer.** Flow cytometric histograms of the nuclear suspension of *C. morifolium* var. 'Sei-marine' and Gojo-0. Genome size of 'Sei-marine' was estimated at 7.93 ± 0.04 Gb ($n$ = 3) according to fluorescence intensity relative to Gojo-0.

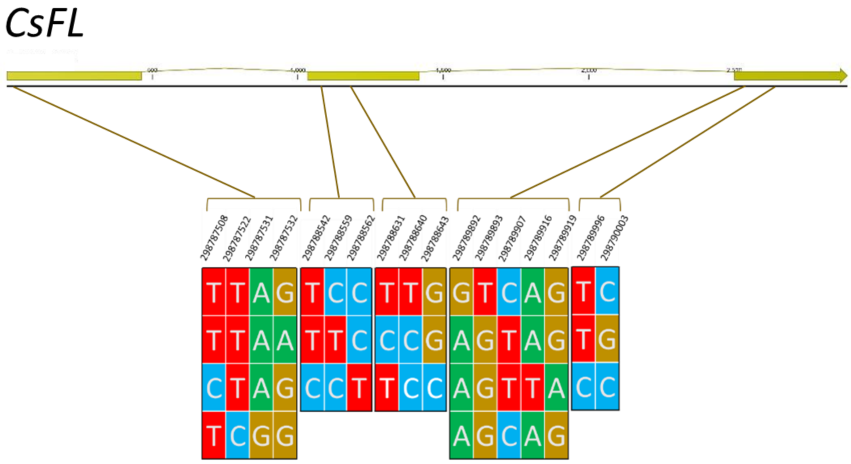


**Supplementary Figure 16 | Haplotype structure of *CsFL* in cultivated chrysanthemum variety 'Jinbudiao'.** Haplotype structures at five sites in *CsFL* based on RNA-seq data of the variety Jinbudiao are shown. The five haplotype blocks are independent.

**Supplementary Table 2 | Classification of intact LTR-RTs in the Gojo-0 genome.**

| Classification | Number |
|---|---|
| **Copia** | **18,820** |
| Ale | 1,676 |
| Angela | 854 |
| Bianca | 21 |
| Ivana | 1,217 |
| SIRE | 5,125 |
| TAR | 1,315 |
| Tork | 267 |
| Others | 8,345 |
| **Gypsy** | **15,085** |
| chromovirus | 3,955 |
| CRM | 153 |
| Galadriel | 72 |
| Reina | 255 |
| Tekay | 2,109 |
| non-chromovirus | 10,251 |
| Athila | 8,512 |
| Tat\|Retand | 1,732 |
| Others | 879 |
| **unknown** | **23,446** |
| **Total** | **57,351** |

**Supplementary Table 4 | Classification of SbdRT copies.**

| SbdRT | | Angela-type | |
|---|---|---|---|
| SbdRT-orf | 161 | 150 | |
| group1 | 46 | 45 | autonomous |
| group2 | 15 | 14 | autonomous |
| others | 6 | 6 | autonomous |
| fragmented | 94 | 85 | nonautonomous |
| SbdRT-nis | 190 | 0 | nonautonomous |
| SbdRT (others) | 9 | 6 | nonautonomous |
| Total | 360 | 156 | |

SbdRT copies with long terminal repeats (LTRs) at both ends are shown. The total number of SbdRT was 360. Of the 161 SbdRT-orf copies, 67 were thought to be autonomous, and 150 were assigned to Angela-type LTR-RT.

**Supplementary Table 9 | Accession numbers of raw data described in this study.**

| Platform | | Library | Bioproject | Accessions | Description | Insert size | Total reads | Total bases |
|---|---|---|---|---|---|---|---|---|
| PacBio | Sequel | WGS | PRJDB7468 | DRX233469 | Single | 13,744bp | 24.9M | 343Gb |
| Illumina | Hiseq2500 | WGS | PRJDB7468 | DRX142424 | PE250 | 619bp | 1,269M | 317Gb |
| Illumina | HiseqX | Hi-C | PRJDB7468 | DRX256944 | PE150 | -- | 1,586M | 238Gb |
| PacBio | Sequel | Iso-seq | PRJDB5536 | DRX274173 | Single | -- | -- | -- |
| | | | | DRX274174 | Single | -- | -- | -- |
| | | | | DRX274175 | Single | -- | -- | -- |
| | | | | DRX274176 | Single | -- | -- | -- |

1. van Geest, G. et al. An ultra-dense integrated linkage map for hexaploid chrysanthemum enables multi-allelic QTL analysis. *Theor. Appl. Genet.* **130**, 2527–2541 (2017).

2. Oda, A. et al. *CsFTL3*, a chrysanthemum *FLOWERING LOCUS T-like* gene, is a key regulator of photoperiodic flowering in chrysanthemums. *J. Exp. Bot.* **63**, 1461–1477 (2012)

3. Shchennikova, A. V., Shulga, O. A., Immink, R., Skryabin, K. G. & Angenent, G. C. Identification and characterization of four chrysanthemum MADS-box genes, belonging to the Apetala1/Fruitfull and Sepallata3 subfamilies. *Plant Physiol.* **134**, 1632–1641 (2004)

4. Higuchi, Y. et al. The gated induction system of a systemic floral inhibitor, antiflorigen, determines obligate short-day flowering in chrysanthemums. *Proc. Natl. Acad. Sci. U. S. A.* **110**, 17137–17142 (2013).

5. Hirakawa, H. et al. De novo whole-genome assembly in Chrysanthemum seticuspe , a model species of chrysanthemums, and its application to genetic and gene discovery analysis. *DNA Res.* **26**, 195–203 (2019).

6. Ma, Y. P., Fang, X. H., Chen, F. & Dai, S. L. DFL, a FLORICAULA/LEAFY homologue gene from Dendranthema lavandulifolium is expressed both in the vegetative and reproductive tissues. *Plant Cell Rep.* **27**, 647–654 (2008).

7. Li, H. et al. The Sequence Alignment/Map format and SAMtools. *Bioinformatics* **25**, 2078–2079 (2009)

8. Huang, D. et al. Identification and characterization of *CYC*-like genes in regulation of ray floret development in *Chrysanthemum morifolium*. *Front. Plant Sci.* **7**, 1–15 (2016).
